# Pattern separation of spiketrains in hippocampal neurons

**DOI:** 10.1101/107706

**Authors:** Antoine D. Madar, Laura A. Ewell, Mathew V. Jones

**Author notes:** contributed equally to the manuscript (Antoine D. Madar, Laura A. Ewell). Author contributions Conceptualization: MVJ, LAE. Data curation: LAE, ADM. Formal analysis: MVJ, ADM. Funding acquisition: MVJ, ADM. Investigation/data collection: LAE, ADM. Methodology: MVJ, LAE, ADM. Project administration: MVJ, LAE, ADM. Resources: N/A. Software: MVJ, ADM. Supervision: MVJ. Validation: MVJ, LAE, ADM. Visualization: MVJ, LAE, ADM. Writing – original draft: ADM. Writing – review & editing: MVJ, LAE, ADM.

## Abstract

Pattern separation is a process that minimizes overlap between patterns of neuronal activity representing similar experiences. Theoretical work suggests that the dentate gyrus (DG) performs this role for memory processing but a direct demonstration is lacking. One limitation is the difficulty to measure DG inputs and outputs simultaneously. To rigorously assess pattern separation by DG circuitry, we used mouse brain slices to stimulate DG afferents and simultaneously record DG granule cells (GCs) and interneurons. Output spiketrains of GCs are more dissimilar than their input spiketrains, demonstrating for the first time temporal pattern separation at the level of single neurons in the DG. Pattern separation is larger in GCs than in fast-spiking interneurons and hilar mossy cells, and is amplified in CA3 pyramidal cells. Analysis of the neural noise and computational modelling suggest that this form of pattern separation is not explained by simple randomness and arises from specific presynaptic dynamics. Overall, by reframing the concept of pattern separation in dynamic terms and by connecting it to the physiology of different types of neurons, our study offers a new window of understanding in how hippocampal networks might support episodic memory.

## Introduction

How does the brain allow us to discriminate between similar events in our past? This question is a central challenge in the neurobiology of memory and remains elusive. To prevent confusion between memories that share similar features, the brain needs to store distinct activity patterns to represent distinct memories. In the influential Hebb-Marr framework of episodic memory^1,2^, representations are stored in area CA3 of hippocampus, an auto-associative network where plastic recurrent excitatory connections facilitate recall of stored patterns in response to partial cues^3,4^. However, strong recurrent excitation severely limits the number of patterns that can be stored without overlap^4,5^. Such overlap would lead, when a partial cue common to several patterns is presented, to the reactivation of many patterns and thus to confusion or confabulation. To avoid these interferences, the Hebb-Marr framework proposes that redundancy between input patterns is reduced before they are stored. This process of transforming similar input patterns into less similar output patterns is termed *pattern separation*^5,6^.

Theoretical models suggest that the dentate gyrus (DG) performs pattern separation of cortical inputs before sending its differentiated outputs to CA3^1,2,4^. Indeed, DG is ideally located to do this, receiving signals via the major projection from entorhinal cortex (EC), the perforant path (PP), and sending signals to CA3 via the axons of granule cells (GCs)^7^. In addition, behavioral studies have shown that DG lesions impair mnemonic discrimination^8-12^ and several experimental reports have shown that similar environments or events are represented differently in the DG^13-17^. However, this separation of DG representations could be inherited from upstream structures (e.g. EC) and simply reported by DG. Therefore, a rigorous demonstration that pattern separation is performed by DG requires simultaneous knowledge of its inputs and outputs^6^. Some electrophysiological studies suggest that EC spatial representations are on average more correlated than in DG^13,14,18,19^, but the recorded EC neurons were unlikely to contact the recorded DG neurons, and were not recorded at the same time: a direct test of whether DG itself performs pattern separation on EC inputs is thus still lacking.

Another difficulty in studying pattern separation is in defining the nature of “activity patterns”. Previous studies have focused on spatial patterns of “active neurons”, with little reference to the dynamics of neural activity. For example, computational models predict that DG separates overlapping populations of active EC neurons into less overlapping populations of active GCs^5,20-23^. Immediate-early genes (IEG) expression studies have confirmed that distinct events drive plasticity in different populations of GCs^15,24^ and that overlap in these representations causes mnemonic interference^25^. In contrast, *in vivo* single-unit recordings in the DG found that similar contexts are represented by the same population of active neurons, but differences are encoded by different spatially tuned firing patterns^13,14,17^.

These conflicting results show that pattern separation could correspond to different computations depending on the type of patterns investigated, and that multiple forms of pattern separation could in theory be implemented by the DG^6^. For example, because *in vivo* recordings suggest that the same neurons can be used to code different environments^13,14,17^, it is possible that pattern separation is performed at the level of single GCs, each disambiguating the activity patterns that it receives. Such disambiguation could be done by changing firing rates, or alternatively, by changing spike timing. Previous experimental investigations of pattern separation in DG examined population vectors of place fields averaged over minutes^13,14,26^, but place cells also carry information at shorter timescales^27-30^. So far, pattern separation has not been well characterized on the scale of milliseconds, and never where patterns are explicitly afferent and efferent trains of action potentials.

Thus, whether the DG network per se reduces the overlap between similar inputs, and how it performs this computation remains a mystery, especially at short timescales. Here, we set out to test the hypothesis that the DG performs pattern separation of cortical spiketrains at the level of single GCs. We designed a novel pattern separation assay in acute brain slices to take advantage of the experimental control afforded to slice electrophysiology. Complex input spiketrains of varying similarities were fed into the DG via its afferents, and the output of a single GC was simultaneously recorded, allowing the first direct measure of pattern separation (by comparing input similarity versus output similarity), on timescales relevant to neuronal encoding and synaptic plasticity. Finally, we explored whether other cell types in the DG and CA3 exhibited this form of pattern separation and investigated the role of neuronal noise and synaptic dynamics in supporting this computation.

## Results

### Temporal pattern separation by individual dentate granule cells

A direct test of pattern separation in single GCs requires knowledge of the similarity between input patterns arriving via the PP and comparison with the similarity between GC output patterns. Here, we define input and output patterns as rasters of spiketrains. The similarity between two spiketrains was assessed by computing their pairwise Pearson’s correlation coefficient (R) using a binning window τ_w_ of 10 ms (unless otherwise specified). We generated sets of Poisson input spiketrains (simulating trains of incoming cortical action potentials), with each set having an average correlation R_input_ (**Figure 1a** and **Methods – Pattern separation experiments**). We then recorded the spiking responses of GCs to these sets of input trains delivered to lateral PP fibers (**Figure 1b-c**) (102 recording sets from 28 GCs in adolescent mice), allowing us to compute the average output correlation (R_output_) (**Figure 2a-b**).

**Figure 1.**
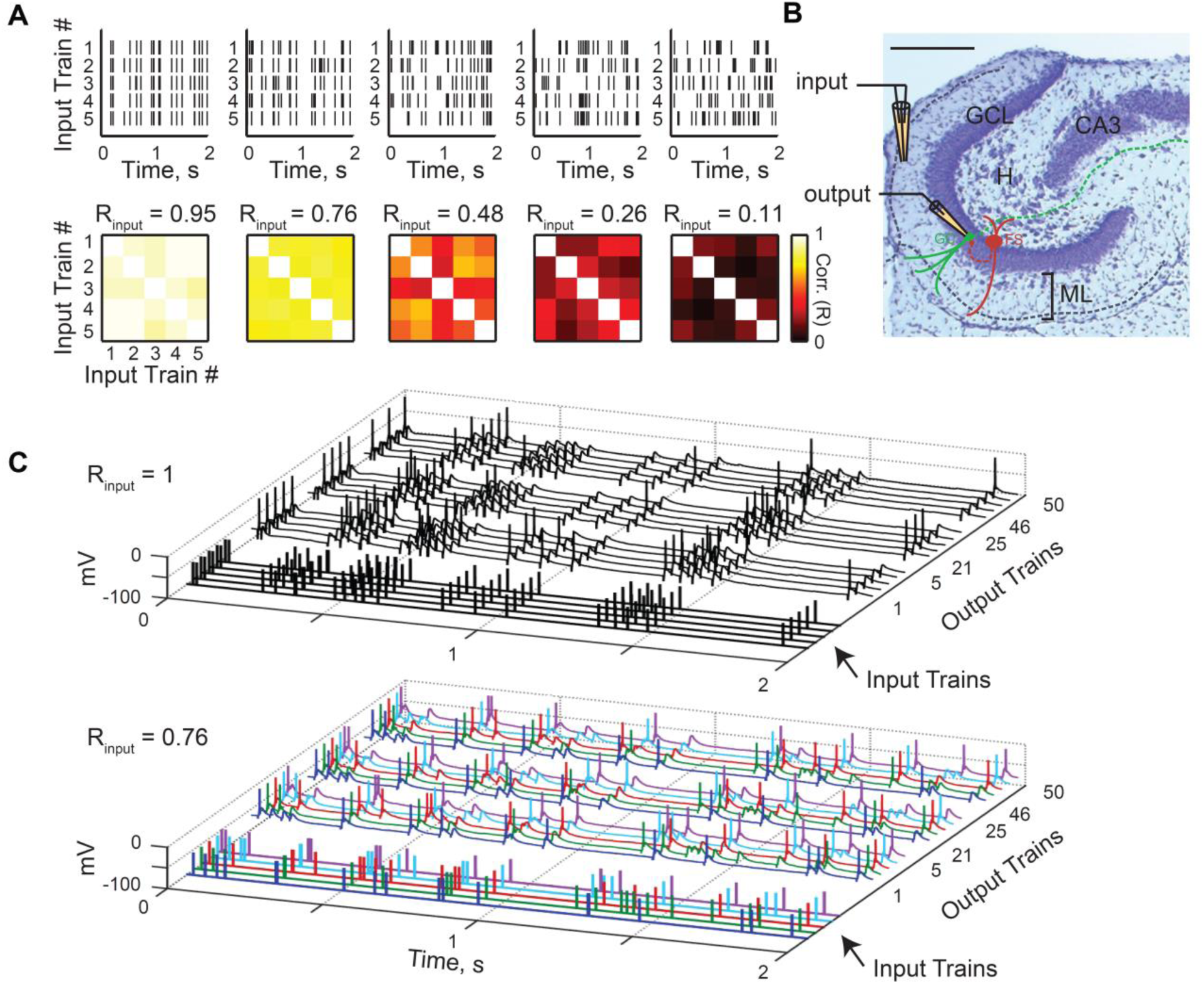
Pattern separation assay in acute brain slices at the single cell level. **(a)** Examples of input sets. *Top*: each input set comprises five different trains of electrical pulses. *Bottom*: correlation coefficient matrix for each input set, each square representing the correlation coefficient between two input trains measured with a binning window (τ_w_) of 10 ms. R_input_ is the average of coefficients, diagonal excluded. **(b)** Histology of the DG in a horizontal slice (Cresyl violet/Nissl staining; scale bar: 250 µm), overlaid with a schematic of the experimental setup: a theta pipette in the ML (input) is used to focally stimulate the PP while a responding GC (output) is recorded via whole-cell patch-clamp. (GCL: granule cell layer, H: hilus, ML: molecular layer, FS: fast-spiking interneuron. Solid lines represent dendrites and dashed lines axons) **(c)** Current-clamp recordings of the membrane potential of two different GCs in response to different input sets (*Top*: R_input_ = 1; *Bottom*: R_input_ = 0.76). Each input set (five input traces) is repeated ten times (only three repetitions are shown, with spikes truncated at 0 mV). In the bottom graph, input trains and their respective children output spiketrains have matching colors.

**Figure 2.**
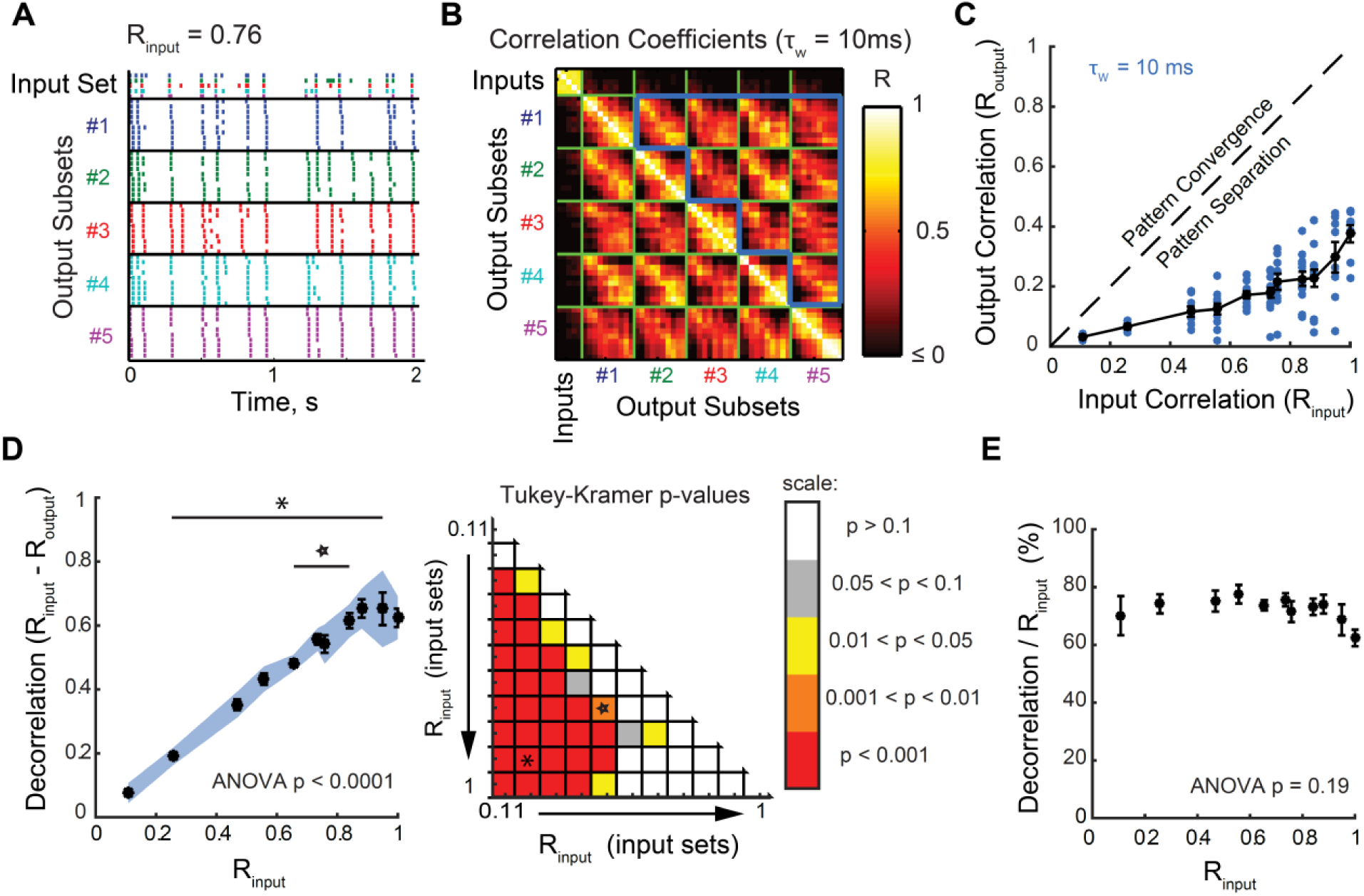
Input spiketrains are decorrelated at the level of individual granule cells. **(a)** Example of a recording set (input set + output set): the raster plot shows one set of input spiketrains and the children output spiketrains recorded from one GC, reordered to display output subsets (i.e., the ten children coming from one parent input spiketrain) together and with the same color. **(b)** Corresponding 55×55 correlation coefficient matrix using a binning window (τ_w_) of 10 ms. Each small square represents the correlation coefficient between two spiketrains. R_output_ is defined as the mean of correlations between individual output spiketrains driven by different input spiketrains, as outlined by the bold blue border, which excludes comparisons between outputs generated from the same parent input. **(c)** Data points, corresponding to 102 recording sets (28 GCs), are all below the identity line (dashed line). This means that R_output_ was lower than R_input._for all recordings, thus demonstrating pattern separation. **(d)** *Left*: Effective decorrelation averaged over all recording sets as a function of R_input_. Although there is a significant decorrelation for all tested input sets (one-sample T-tests: the blue shade indicates the 95% confidence interval that average decorrelation is significantly above 0), they are effectively decorrelated to different magnitudes (one-way ANOVA, p < 0.0001). *Right:* Matrix of p-values from post-hoc Tukey-Kramer tests comparing effective decorrelation levels across all pairs of parent input sets. The asterisk and star correspond to the comparisons displayed in the left panel. This analysis shows that the decorrelation is significantly different (higher) for highly similar input spiketrains than for already dissimilar inputs. **(e)** When the effective decorrelation is normalized to the correlation of the input set, there is no significant difference between input sets (ANOVA, p = 0.18). In all graphs, τ_w_ = 10 ms. Means and SEM in black.

For every recording set, R_output_ was lower than the R_input_ of the associated input set, indicating a decorrelation of the output spiketrains compared to their inputs (**Figure 2c**). This was also the case in GCs recorded in slices from adult mice (35 recording sets from 14 neurons) (**Figure 9d**). The effective decorrelation, defined as the difference between R_input_ and R_output_, was statistically significant for every input set (**Figure 2d left**), but was larger when input spiketrains were highly correlated (**Figure 2d**). This is consistent with the role of DG in discriminating between similar memories more than already dissimilar ones ^9^. Note, however, that the decorrelation normalized to R_input_ is invariant: whatever the input set, the output trains were always decorrelated to ∼70% of R_input_ (**Figure 2e**). Such invariance suggests that the same decorrelating mechanism is used on all input sets.

These results constitute the first direct experimental demonstration that single GCs, the output neurons of DG, exhibit temporal pattern separation. A comparison of the spiking patterns recorded in multiple GCs also shows that different GCs tend to process the same input spiketrain in different ways (**Figure S1**). Such diversity could support pattern separation at the population level.

### Temporal pattern separation in other cell types of the DG network

Any channel processing inputs and returning outputs, and thus any brain network, performs either pattern separation or pattern convergence to some degree^6^. Thus, GCs are unlikely to be the only neurons to exhibit temporal pattern separation of spiketrains. However, we would expect pattern separation to be at its greatest in GCs, at least among DG cells, because they are the output neurons of the DG and thus should provide the most separated patterns to CA3 before they are stored. To test this hypothesis, we performed the same pattern separation assay while recording from fast-spiking interneurons (FS, 20 recording sets) (**Figure 3a**) or hilar mossy cells (HMC, 18 recording sets) (**Figure 3b**).

**Figure 3.**
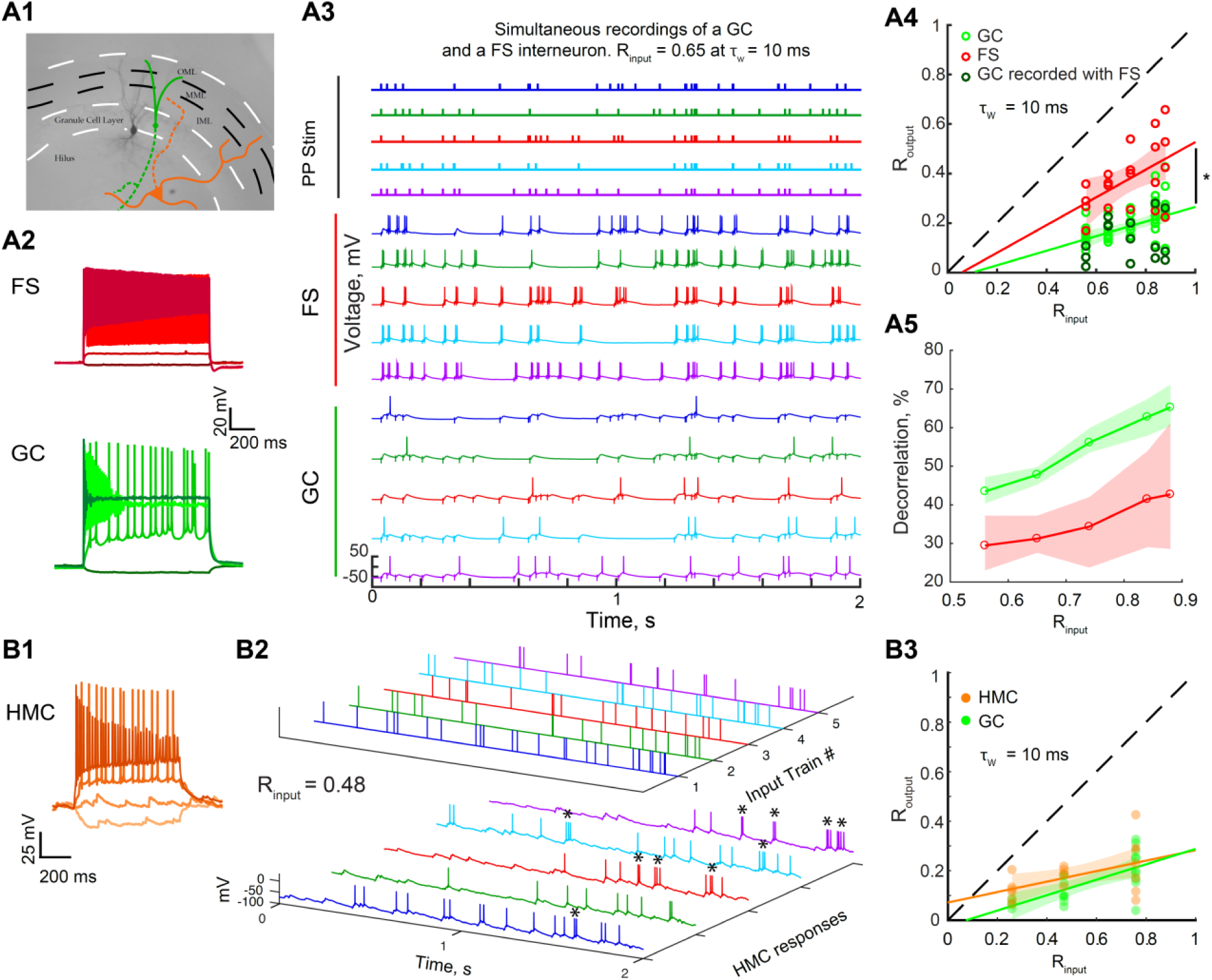
Different levels of temporal pattern separation in different DG cell types. **(a1)** Picture of a recorded FS filled with biocytin (black). In the case of simultaneous recordings, the recorded GCs were close to the FS, as depicted by the schematic in green. In different experiments, we recorded from putative hilar mossy cells (HMC, orange). Full lines represent dendrites, dashed lines axons. **(a2-3)** Example of a simultaneous whole-cell recording of a GC and a neighboring FS. **(a2)** Simultaneous membrane potential recordings (baseline at −60mV) of a FS and a GC to the same set of current steps (−25, 100, 500 and 1000 pA). **(a3)** Simultaneous current-clamp recordings of the same FS and GC as in a2 in response to the five input traces of an input set with R_input_ = 0.65. Simultaneous input and output trains have the same color. **(a4)** R_output_ versus R_input_ for FSs and GCs. Data points correspond to recording sets: 20 for FS (red, 4 cells, 4 per input set), and 61 for GC (green, with a darker shade open circle when simultaneously recorded with a FS, 13 cells, 11-13 per input set). All GC recordings done at the same input correlations as FS recordings were used for an unpaired comparison showing that FS exhibit less spiketrain decorrelation than GCs: ANCOVA: p < 0.0001, 95% confidence bounds around the linear fits shown as shaded areas; two-way ANOVA: input sets: p = 0.0016, cell-types: p < 0.0001, interaction: p = 0.72. Post-hoc T-tests with Bonferroni corrections for 5 comparison groups: all p < 0.05 (for decreasing R_input_, p = 0.0307, 0.0181, 0.0007, 0.0001, 0.0122) **(a5)** Effective decorrelation (R_input_ – R_output_) for FS and GC. Shaded areas represent the 95% confidence interval of a bootstrap test comparing the mean decorrelation of both celltypes: GCs exhibit significantly more pattern separation than FS for all input sets. (**a4-5**) Note that when comparing only the simultaneous GC and FS recordings, we found a similarly significant difference between celltypes. (**b1**) Membrane potential of a hilar neuron in response to current steps (−100, 0, 100, 400 pA; baseline at −70mV), showing a spontaneous barrage of EPSPs, regular spiking, and a lack of large after-hyperpolarization, all typical features of HMCs (Larimer & Strowbridge, 2008). (**b2**) Current-clamp recordings of same HMC in response to a set of five input trains (R_input_ = 0.49, τ_w_ = 10ms). HMCs fire occasional bursts of spikes (marked by asterisks) in response to a single input, which was not seen in GCs. (**b3**) R_output_ versus R_input_ for HMCs and GCs. Data points correspond to recording sets: 18 for HMC (orange, 11 cells, 5-7 per input set), and 22 for GC (green, 11 cells, 4-10 per input set). An unpaired comparison suggests that HMCs and GCs show only slight differences in pattern separation measured at τ_w_ = 10 ms: ANCOVA: p = 0.15, 95% confidence bounds around the linear fits shown as shaded areas; two-way ANOVA: input sets: p = 0.0004, cell-types: p = 0.074, interaction: p = 0.57. Post-hoc T-tests with Bonferroni corrections for three comparison groups (for decreasing R_input_): p = 1, 0.05, 0.21.

At the 10 ms timescale, the distributions of average (R_input_, R_output_) were significantly different between FSs and GCs, with the R_output_ of simultaneously recorded GCs always lower than their corresponding FS (**Figure 3a4**):FSs exhibit lower levels of decorrelation than GCs (**Figure 3a5**). For HMCs, although their current-clamp responses and spiking behavior looked different from GCs (**Figure 3b2**), the distribution of average (R_input_, R_output_) appeared only slightly different from GCs with this analysis (**Figure 3b3**, but see below, in **Figure 5d**).

### Temporal pattern separation in CA3 pyramidal cells

Our FS recordings have shown that not all neurons necessarily exhibit as high levels of temporal pattern separation as GCs. We thus asked whether this form of pattern separation is specific to the DG output, or whether neurons in other hippocampal regions can perform temporal pattern separation. To answer this question, we adapted our pattern separation assay to CA3 pyramidal cells (PCs) (**Methods – Pattern separation experiments**) and recorded their output spiketrains in response to direct stimulation of the GCL, i.e. in response to its input from the DG (**Figure 4a**). Due to strong feedforward inhibition, CA3 PCs do not generally spike in response to external stimulation of afferents in slices, unless inhibition is totally blocked^31-33^. By using 30 Hz Poisson input sets (**Figure 4b**) and adding 100 nM of gabazine to the bath, which only slightly decreases the amplitude of GABA-A-mediated IPSCs (see **Methods**), we managed to record for the first time the spiking output of CA3 PCs in response to complex input spiketrains while preserving some inhibition in the network (**Figure 4c**). PCs fired during periods of high input frequency, probably due to depression of local inhibitory transmission^31,34^ combined with facilitation at the mossy fiber-CA3 PCs synapses^35^, and as a result action potentials often appeared more clustered than in control GC recordings (**Figure 4c**). Despite this spiking pattern, we found that, at the 10 ms timescale, CA3 PCs average (R_input_, R_output_) distribution was slightly but significantly lower than for GCs recorded in similar pharmacological conditions (**Figure 4d**). This indicates that CA3 PCs are different from GCs in respect to temporal pattern separation, but that, surprisingly, their output spiketrains are even more decorrelated. Temporal pattern separation is thus not specific to the DG.

**Figure 4.**
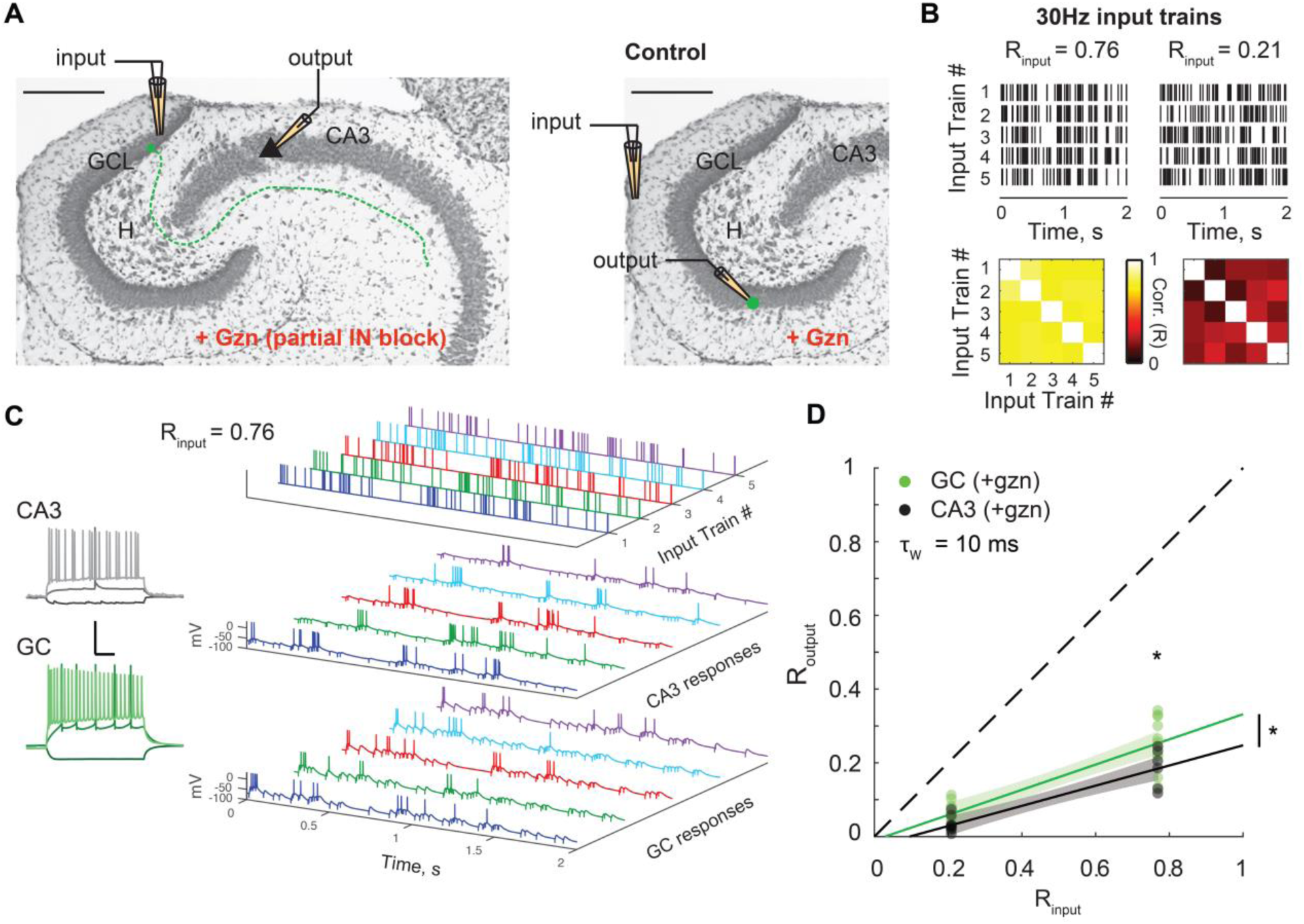
CA3 pyramidal cells exhibit more temporal pattern separation than GCs. **(a)** Schematic of the experimental setup: a theta pipette in the granule cell layer (GCL) is used to stimulate GCs and their mossy fibers (dashed line) to evoke action potentials in CA3 pyramidal cells recorded via whole-cell patch-clamp. To limit feedforward inhibition and allow pyramidal cells to spike, experiments were performed under partial block of inhibition (100 nM of gabazine). Control experiments were performed in GCs under the same pharmacological conditions but with OML stimulations. **(b)** Two input sets of 30 Hz Poisson trains were used. *Top*: rasters of the five spiketrains of each set. *Bottom*: correlation coefficient matrix for each input set (τ_w_ = 10 ms). **(c)** Example of current-clamp recordings from a pyramidal cell and a GC from the same animal. *Left:* Membrane potential responses to current steps (−100pA, 100pA, 350pA). Scale: 100 ms (horizontal) and 50 mV (vertical). *Right:* example of 5 output responses (sweeps 11-15) to an input set (R_input_ = 0.76). **(d)** R_output_ versus R_input_ for CA3 pyramidal cells and GCs. Data points correspond to recording sets: 15 for CA3 (black, 14 cells, 6-9 recordings per input set), and 22 for GC (green, 13 cells, 11 per input set). The CA3 distribution is lower and significantly different from the GC distribution: ANCOVA: p < 0.01, 95% confidence bounds around the linear fits shown as shaded areas; two-way ANOVA: input sets: p < 0.0001, cell-types: p = 0.0036, interaction: p = 0.24. Post-hoc T-tests with Bonferroni corrections for two comparison groups: p = 0.032 for R_input_ = 0.76, p = 0.1 for R_input_ = 0.21.

**Figure 5.**
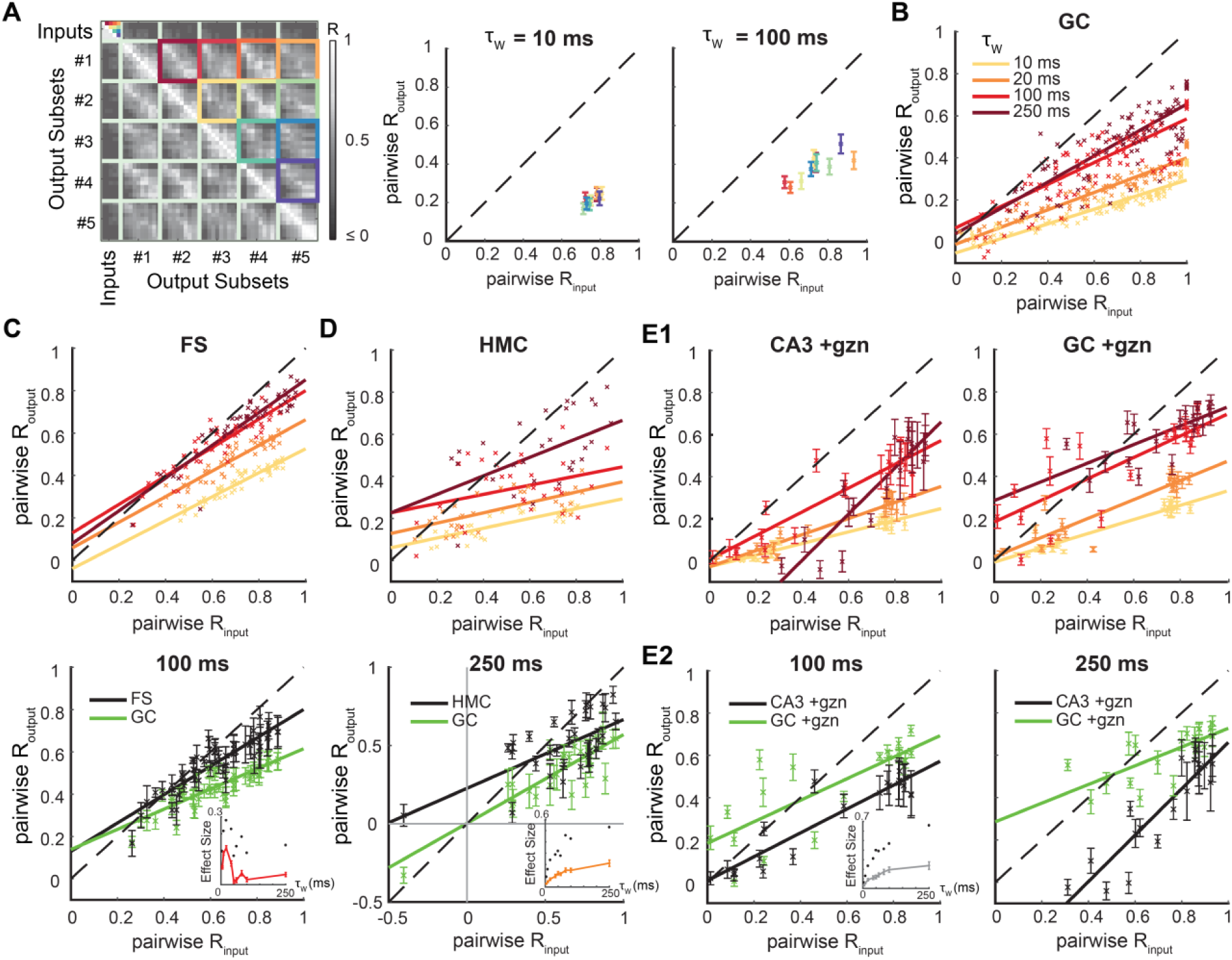
Differences in temporal pattern separation between hippocampal celltypes depend on the timescale. **(a)** Pairwise analysis. Instead of averaging across all five input trains of a set (here R_input_ = 0.76 at τ_w_ = 10 ms, same matrix as in Fig. 2b), we average children output coefficients corresponding to a single pair of input trains (identified by color-coded squares.). This finer analysis is necessary at τ_w_ higher than 20 ms because the *pairwise input correlation coefficients* are not well constrained around their mean anymore (see right panel). *Middle* and *Right*: mean across cells, following the same color-code as displayed in the matrix (*Left*). **(b)** Pairwise R_output_ vs pairwise R_input_ for GCs, measured with different binning windows τ_w_. Only the means across cells are displayed, but the regression line was fitted on the full distribution of data points. The larger τ_w_, the less GCs exhibit pattern separation. **(c)** *Top:* Pairwise analysis on FS recordings (same as in b). *Bottom*: mean± SEM and regression lines for τ_w_ = 100 ms: FS and GC distributions are still different, especially at high input similarity (ANCOVA: p < 0.0001 for τ_w_ = 10 up to 250 ms). Lower right inset: effect size as a function of the timescale τ_w_ (up to 250 ms): Mean ± SEM (red) and maximum (black dots) of the absolute difference between FSs and GCs Routput mean values for all pairwise R_input_. Note a decreasing effect of larger τ_w_ on the difference between FSs and GCs in terms of pattern separation. **(d)** *Top:* Pairwise analysis on HMC recordings (same color code as in b). *Bottom*: At τ_w_ = 250 ms, HMC and GC distributions are different (ANCOVA: p < 0.0001 for τ_w_ = 10 up to 250 ms), especially at lower input similarity: notice the points above the identity line, showing pattern convergence for HMCs in contrast to GCs. Under this pairwise analysis (in contrast to Fig. 3b), HMCs and GCs are significantly different at τ_w_ = 10 ms (comparison graph not shown, see inset for effect size). However, the inset (see c for legends) shows an increasing effect of τ_w_ on the difference between HMCs and GCs in terms of pattern separation. (**e1**) Pairwise analysis on CA3 pyramidal cells and GCs, both under partial inhibition block and under 30 Hz inputs (same color code as in b). (**e2**) At τ_w_ = 100 ms and 250 ms, CA3 and GC distributions are still different (ANCOVA: p < 0.0001). The inset (see c for legends) shows an increasing effect of τ_w_ (plateauing at large timescales) on the difference between CA3 PCs and GCs.

### Temporal pattern separation at different timescales across celltypes

Because the timescales meaningful for the brain remain uncertain, it is important to assess the separation of spiketrains at different timescales. We therefore binned spiketrains using a range of τ_w_ from 5 ms to 250 ms and performed a finer grained analysis using pairwise R_input_ and associated pairwise R_output_ instead of the average across the ten pairs of input trains (**Figure 5a**). We discovered that pattern separation levels can dramatically change as a function of τ_w_. In GCs, the larger the timescale the less they exhibit decorrelation of their input spiketrains, especially at high R_input_ (**Figure 5b**). Nonetheless, GCs still exhibit relatively high levels of pattern separation of highly similar input spiketrains, even at long timescales (**Figure 5b**).

This analysis confirmed that, at short timescales, FSs exhibit less pattern separation than GCs (**Figure 5c**) and revealed significant differences between HMCs and GCs, especially for pairs of input spiketrains with low R_input_ (**Figure 5d**), which were not as obvious in our previous coarse analysis (**Figure 3b3**). At longer timescales, the variability across neurons has a tendency to increase for all celltypes, but the average levels of decorrelation of both FSs and HMCs stayed highly significantly different from those of GCs (**Figure 5c-d**). Interestingly, the difference between FSs and GCs was larger at short timescales, whereas the difference between HMCs and GCs increased with larger timescales. Indeed, at τ_w_ above 100 ms, HMCs can often exhibit pattern convergence instead of pattern separation, especially for pairs of already dissimilar input spiketrains (low or negative pairwise R_input_), whereas FSs just show weak or no pattern separation.

Concerning CA3 PCs, they still exhibit high levels of temporal pattern separation at long timescales and the difference with GCs increases (**Figure 5e**). Interestingly, in contrast to all other tested celltypes and conditions (**Figure 5b-d**), PCs even show a dramatic increase of their average levels of decorrelation at the 250 ms timescale (**Figure 5e**).

Overall, these results show that among the tested DG celltypes, GCs exhibit the highest levels of temporal pattern separation of cortical spiketrains across all timescales. Our findings also suggest that the high level of separation by the DG is amplified in CA3.

### Mechanism of temporal pattern separation: neural noise

To determine what mechanisms might support temporal pattern separation in GCs, it is necessary to understand its dynamics first. Limiting our analysis to the first presentation of an input set revealed that outputs were already significantly decorrelated (**Figure 6a-b**). This shows that the separation mechanism is fast, consistent with the fact that the brain generally does not have the opportunity to average repeated signals and that separation must happen immediately during encoding. In addition, analysis of the last presentation revealed only modestly more separation than for the first one, and only for high input correlations (**Figure 6c**), suggesting that learning to recognize the input pattern is not critical.

**Figure 6.**
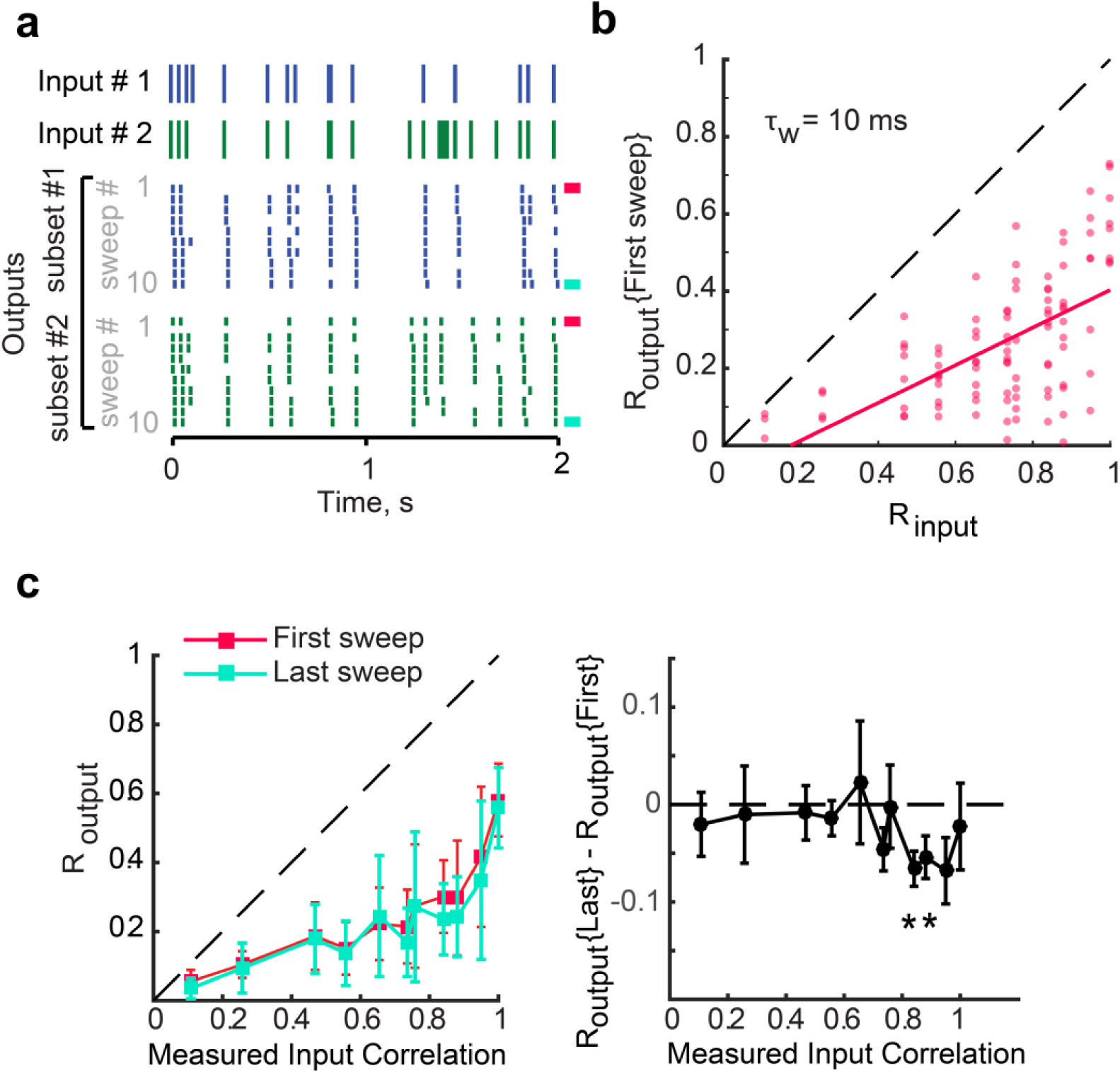
Input spiketrains are efficiently separated upon their first presentations. **(a)** Two of five inputs are shown with corresponding output spiketrains. The first output sweep is marked with a pink bar (right) and last sweep is marked with a blue bar. **(b)** R_output_, computed from the first sweep of five output trains only (pink), as a function of R_input_, fitted with a parabola. All data points are below the identity line indicating that outputs are effectively decorrelated compared to their inputs even when input patterns have only been presented once each. The average decorrelation (R_input_-R_output_) is significant for all input sets (one-sample T-tests, p < 0.01) except for R_input_ = 0.11 (p = 0.1). **(c)** *Left*: Output correlations (Mean ± SEM) between spiketrains of the first sweep (pink) and the last sweep (blue). There is no significant difference (ANCOVA, p = 0.33). *Right:* When taking into account that the two distributions are paired, we detect that a few output correlations are significantly lower for the last sweep than for the first one (one-sample T-test on the difference between R_output_ of the first and last sweep of each recording set, asterisks signify p < 0.05). This is evidence, though weak, that repetition of input spiketrains might improve pattern separation for highly similar inputs.

Because the mechanism for temporal pattern separation is fast and does not require learning, we asked first whether intrinsic properties of GCs could play a role. Linear regression analysis revealed that the membrane capacitance, resistance, time constant as well as the resting membrane potential are not predictors of decorrelation in GCs (see low R^2^ in **Table S1**). Another hypothesis is that randomness in neuronal responses drives the decorrelation. Indeed, when the same input spiketrain is repeated (e.g. R_input_ = 1) the output spiketrains are not well correlated (as shown by the mean spiketrain reliability R_w_) (**Figure 7a**), consistent with well-known trial-to-trial variability in single neuron responses^36-38^. Theoretical investigation of pattern separation often relies on some sort of random process such as probabilistic neuronal activation^5^ or stochastic firing^39^, which suggests that “neural noise” is a likely contributor to any form of pattern separation. However, because “neural noise” can cover multiple different definitions and phenomena^36^, determining its role in a complex computation is not trivial.

**Figure 7.**
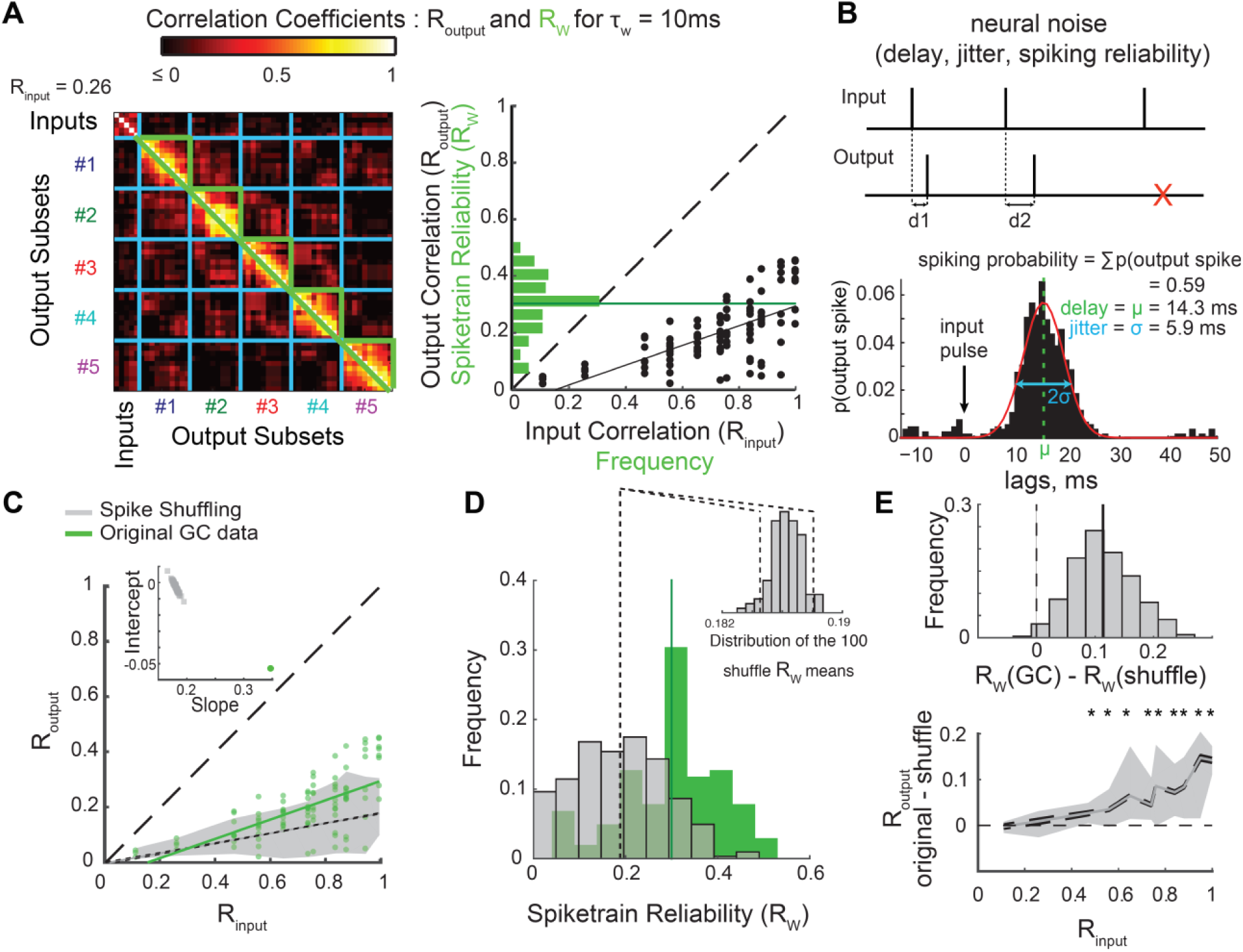
Pattern separation in single GCs is not explained by simple neural noise. **(a)** The variability of output spiketrains in response to the same input train sets the upper bound for R_output_. *Left:* Correlations between pairs of output spiketrains associated with different input trains (R_output_) and pairs of different output spiketrains associated with the same input train (enclosed by green, R_w_: spiketrain reliability, the reproducibility of the output given the same input). *Right:* Frequency distribution of R_w_ for all recordings (green; dark green line is the mean: <R_w_> = 0.3), overlaid on the distribution of 102 (R_output_, R_input_) points and its regression line (black). Note that means <R_w_> and <R_output_> for R_input_ = 1 are close because they both assess the reproducibility of the output when the input is the same. **(b)** Characterization of neural noise. *Top:* example of input and output spiketrains illustrating variable delay of the response spike after an input spike (d1, d2) or failure to spike after an input spike (red cross). *Bottom*: Example from one GC recording. The spike-wise noise in output spiketrains is characterized by the average spike delay, the standard deviation of this delay (jitter) and the probability of spiking after an input spike (spiking probability, SP). **(c-e)** Effect of spike shuffling on R_output_ and R_w_. For each output spiketrain, each original spike was reassigned to the time of a randomly chosen input spike. This shuffling was performed 100 times for each of the 102 original GC recording sets, producing 10,200 shuffled recording sets. **(c)** The (R_output_, R_input_) GC distribution (green) is overlaid with the 90% sample interval of the distribution of the 10,200 shuffled recording sets (grey area: 5 to 95 percentiles). A linear regression was performed for each of the 100 shuffling distributions (102 simulated data points per shuffle): the 90% confidence interval (CI) around the mean regression line is represented by the two dashed lines (short dash) very close to each other, and the GC regression line (green) falls out of this range. As illustrated in the inset, 100% of the shuffling regression lines (grey squares) have a lower slope and higher intercept than the regression line for the original GC dataset (i.e. the green dot is outside of the grey cloud). This Monte-Carlo exact test shows that spiketrains are significantly *less* separated in GCs than would be expected from random spiking. **(d)** Frequency distributions of R_w_ for 102 GC recording sets (green, same as in a, solid line is mean) and 10,200 shuffled recording sets (grey). Dashed lines represent the 90% CI of the shuffling mean (see inset). Inset: distribution of the mean R_w_ for the 100 shufflings. GCs mean R_w_ (0.3) is outside of this distribution, showing that GCs output is significantly *more* reliable than expected from random spiking. **(e)** Paired statistical tests (based on difference between each GC recording and its 100 associated shuffled recording sets) show that spike shuffling leads to smaller R_output_ and R_w_ than original GC recordings. *Top:* Frequency distribution of the difference of R_w_ (10,200 data points). Monte-Carlo exact tests: 99.25% of data points (i.e. p = 0.0075) and 100% of the shuffling means are above 0. *Bottom*: Difference of R_output_ as a function of R_input_. The grey area represents the sample interval where 90% of the 10,200 data points fall (5-95 percentiles): for high R_input_ values, all points are above 0 (see table S4 for details). Solid grey line: means; dashed lines: 90% CI. Monte-Carlo exact tests based on proportion of shuffling means above 0: asterisks denote significance (see table S4 for details).

The noisiness in neural communication is often understood as the unreliability of spiking after a single input spike, and the jitter of the delay between an input spike and an output spike^40,41^. We thus assessed such spike-wise noise directly from recording sets from our pattern separation experiments (**Figure 7b**, **S2**, and **Materials and methods – Analysis of output spiketrains**). Although the spike probability (SP), delay and jitter of GCs were slightly higher in our experiments than in a previous report (but recording and analysis methods were different), the variability between GCs was consistent^41^. We then asked whether these spike-wise noise parameters could predict the degree of decorrelation by GCs. First, linear regression analysis shows no good relationship with any parameter, the SP being an average predictor at best (**Figure S3a** and **Table S2**). Moreover, the average firing rate of a GC output set (a measure dependent on SP) is not well correlated with the degree of decorrelation either (**Table S3** and **Figure S8a**). Temporal pattern separation in GCs seems to not be achieved merely because their output spiketrains are a sparser and jittered version of their inputs.

To more carefully test the hypothesis that random spiking failures and random delays support fast temporal pattern separation, we simulated recording sets based on computational models only governed by spike-wise noise statistics comparable to the original data (**Figure S2, S3b, Table S2** and M**ethods – Computational models**). The distribution of (R_input_, R_output_) was significantly higher in the original data (**Figure 7** and **S5**), showing that simple random processes yield greater levels of separation than real GCs, especially for highly similar inputs (**Figure 7e** and **S5a)**.

In addition to the spike-wise noise, we considered neural noise at the level of spiketrains by computing the average correlation coefficient *R*_*w*_ between “children” output spiketrains from the same “parent” input train (**Figure 7a**). *R*_*w*_ characterizes the more complex notion of *spiketrain reliability*, that is the ability of a neuron to reproduce the same output spiketrain in response to repetitions of the same input spiketrain. *R*_*w*_ is not dependent on intrinsic cellular properties (**Table S1**) and only moderately determined by spike-wise noise parameters (**Table S2**), suggesting that the rather low R_w_ of GCs is the expression of more complex noisy biophysical processes. Consistently, R_w_ was significantly lower for shuffled and simulated data than in real GCs (**Figure 7d-e** and **S5b**). This indicates that the output spiketrains of GCs are more reliable than if their output was entirely determined by simple random processes. Overall, GCs have both a higher R_output_ and R_w_ distributions than datasets based on random spiking (**Figure 7**), which clearly shows that simple noise cannot fully underlie the operations performed by GCs on input spiketrains.

The fact that random spiking yields better temporal pattern separation but less reliable information transmission than GCs also suggests that there might be an unavoidable trade-off between achieving pattern separation and reliable information transmission about input spiketrains. To further investigate this, we looked at the relationship between the spiketrain reliability R_w_ and the decorrelation levels in GC recordings and found a strong anticorrelation (**Figure 8a** and **Table S3**). This linear relationship is clear evidence that biological processes leading to sweep-to-sweep variability is a powerful mechanism for temporal pattern separation in DG.

**Figure 8.**
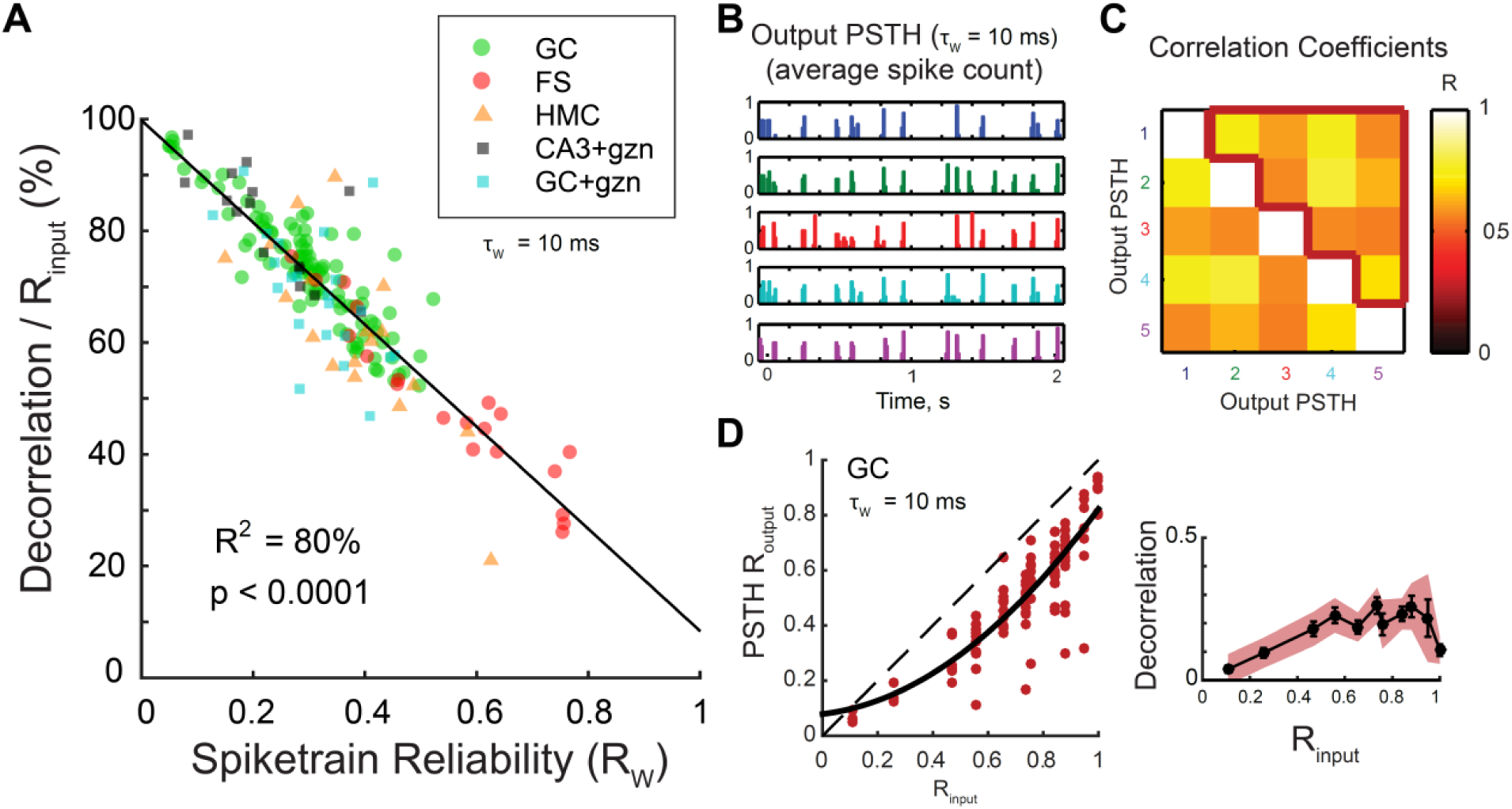
Unreliability in spiketrain transmission is a major but not unique source of temporal pattern separation. **(a)** Spiketrain reliability (R_W_) is an excellent predictor of normalized decorrelation (defined in Figure 2E) for all celltypes and conditions. Notice that, despite the strong anticorrelation, the intercept of the linear model at R_w_ = 1 predicts that even a perfect reliability could still allow 10% of decorrelation. See Table S3 for linear regressions on single celltypes. **(b)** To assess the amount of decorrelation not due to spiketrain unreliability, the ten children output spiketrains of each of the five trains of an input set can be averaged to give the five output peristimulus histograms (PSTH). The 10 ms binned PSTHs of the output rasters in Figure 2a are shown. **(c)** Correlation coefficients between all pairs of the five output PSTHs. The mean correlation (PSTH R_output_) is the average of coefficients inside the red border, and excludes self-comparisons. **(d)** *Left:* PSTH R_output_ as a function of R_input_ (102 recording sets, in red), fitted with a parabola (black). All points are below the identity line indicating decorrelation of outputs compared to inputs. *Right:* Average effective decorrelation (R_input_ – PSTH R_output_) as a function of R_input_ (bars are SEM) reveals a significant decorrelation for all input sets except for the most dissimilar (one-sample T-tests; shaded area is the 95% confidence interval for significant decorrelation).

**Figure 9.**
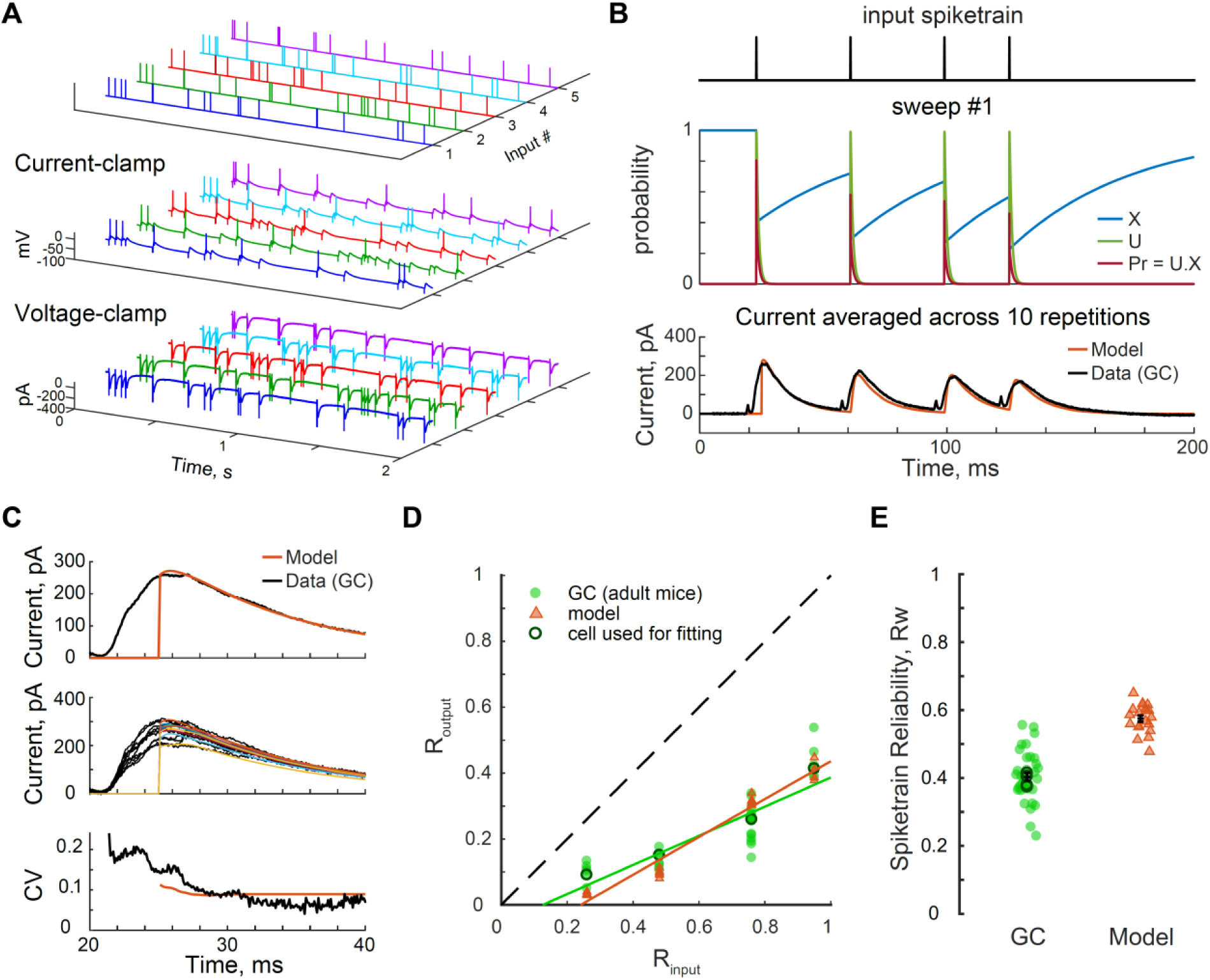
Probabilistic presynaptic release is a potential mechanism balancing temporal pattern separation and spiketrain reliability. **(a)** Non simultaneous current-clamp (V_rest_ ≈ −70mV) and voltage-clamp recordings (V_hold_ = −70mV) in response to a set of input patterns were successively performed in GCs from adult mice. The example shows the first sweep of responses from a single cell to the five trains of an input set (R_input_ = 0.76, τ_w_ = 10 ms). **(b-e)** The recording set of the example cell in a was used to fit a computational model of a spiking neuron with dynamic and probabilistic synapses. **(b)** *Top*: four first input spikes of input train #1. *Middle*: Dynamics of the variables of a probabilistic Tsodyks-Markram model of a synapse in response to the four input spikes. X represents the probability of vesicle availability at the presynaptic site, U represents the probability of release of available vesicles, and Pr is the probability of release for all vesicles. The number of released vesicles was simulated at every time point from a binomial process. *Bottom*: Parameters of the model were adjusted so that the resulting current (averaged over the 10 repetitions, in dark orange) would reasonably match the peak of the corresponding average EPSCs in the original voltage-clamp recordings (baselined and inverted display with partially blanked stimulation artifact, in black). **(c)** Parameters of the model were also adjusted to match the variability in the amplitude of the first EPSC in the original recording. *Top*: average EPSC. *Middle*: individual EPSCs evoked from the 10 repetitions of the first input spike in input train #1. The range of peak amplitudes is similar between model and data. *Bottom*: The coefficient of variation of the current in the model is close to the data. **(d-e)** Temporal pattern separation and spiketrain reliability (τ_w_ = 10 ms) for GCs from adult mice and a single model regular spiking Izhikevich neuron with probabilistic synapses as described above. Data points correspond to recording/simulation sets: 35 for GCs (green circles, 14 cells, 6-16 per input set), and 5 simulations at all input sets (dark orange triangles), with the model fitted to a single recording set (R_input_ = 0.76; R_output_ and R_w_ shown in dark green circles for all input sets).

However, spiketrain unreliability is not the only reason GCs output spiketrains are decorrelated. Indeed, three lines of evidence support the idea that even if GCs were perfectly reliable (i.e. a given input train always leads to the same output train), the set of output spiketrains would still be less similar than the set of inputs. First, the linear model describing the relationship between Rw and decorrelation levels suggest that even at Rw = 1, ∼10% of decorrelation would still be achieved (**Figure 8a**). Moreover, high levels of pattern separation are performed upon the first presentation of an input set (**Figure 6**): this means that even if the full output set was composed of exact repetitions of the set of output trains recorded during the first sweep (e.g. Input trains 1 and 2 always lead to the same Output trains 1 and 2, respectively) pattern separation would still occur, at the levels measured for the first sweep (difference between Outputs 1 and 2 > difference between Inputs 1 and 2). In other words, perfect reliability does not equate the absence of pattern separation. Finally, when averaging out the variability between spiketrains associated to the same input (i.e. by computing the PSTH), a significant level of decorrelation is still detected (**Figure 8b-d**). We can thus conclude that temporal pattern separation is supported by some mechanisms in addition to those that produce spiketrain unreliability.

Taken together, these results suggest that complex but noisy biophysical mechanisms allow GCs to balance temporal pattern separation and reliable signaling about their inputs.

### Mechanism of temporal pattern separation: short-term synaptic dynamics

The fact that intrinsic properties of GCs do not predict their decorrelation levels (**Table S1**) suggests that temporal pattern separation comes from the DG network in which each recorded GC is embedded. We thus hypothesize that the specific short-term dynamics of synapses in the DG network implement temporal pattern separation in GCs. This is a likely mechanism because: 1) Synaptic transmission is a stochastic process and generally considered the main source of noisiness in the spiking output of neurons^36,42^, which would fit our conclusion that temporal pattern separation is dependent on neural noise (see paragraph above), 2) Short-term plasticity makes the probability of synaptic release dependent on the timing of the preceding input spikes, which could explain why temporal pattern separation and spiketrain reliability are not purely governed by noise. To test the idea that short-term synaptic dynamics is a potential mechanism controlling the pattern separation/spiketrain reliability ratio in GCs, we developed a computational model of a spiking GC with dynamic and probabilistic PP-GC synapses (see **Methods – Computational models**). The parameters of the model controlling the facilitation, depression and variability of presynaptic release, and thus the amplitude of EPSCs, were constrained to match the whole-cell voltage-clamp recording of a GC in response to a pattern separation protocol (10 Hz input trains, R_input_ = 0.76) (**Figure 9a-c** and see **Methods – Computational models**). Simulations of pattern separation experiments with this model resulted in high levels of output spiketrains decorrelation, akin to real GCs, but also in relatively high levels of spiketrain reliability (**Figure 9d-e**). This is a proof of principle that the probabilistic and dynamic nature of presynaptic release at the PP-GC synapse is a potential mechanism balancing temporal pattern separation and information transmission in the DG.

### Sources of pattern separation differences between celltypes

To further understand the mechanisms behind temporal pattern separation in single hippocampal cells, we investigated the sources of the difference in decorrelation levels between the tested celltypes. We considered differences in intrinsic cellular properties (**Table S1**), firing rate and probability of bursting (**Figure S7-8**), spike-wise noise (**Figure S3-4**), spiketrain reliability (**Figure 8a** and **S6**) and synaptic transmission dynamics (**Figure S9**).

Both FS and HMC recordings displayed bursts of spikes (defined as more than one output spike between two input spikes), which was very rarely seen in our GC and CA3 recordings (**Figure 3-4, S7c-e,** and **S8c, e**). As a result, both FSs and HMCs had significantly higher firing rates than GCs (**Figure S7a-b**) (although the effect size was smaller for HMCs, because they burst less often than FSs and they have generally less spikes per bursts. **Figure S7d-e**). Then, are the firing rate or the probability of bursting predictive of differences in pattern separation? The relationships are unclear, but it seems that both could partially explain the lower decorrelations observed in some HMCs and FSs (**Figure S8a-b**). To more directly test the role of bursting, we processed all FS and HMC recordings by removing all but the first spike in each detected burst (**Figure S8c, e** and see **Methods – Analysis of the output spiketrains**). These resulting “non-bursty” datasets (nbFS and nbHMC) still exhibited significantly less pattern separation than GCs (and even pattern convergence of dissimilar input trains, in the case of HMCs) (**Figure S8d, f**). Therefore, bursting and high firing rates, although a source of differences in temporal pattern separation, are not a sufficient explanation.

Although FSs show less pattern separation than GCs, it is interesting that they do exhibit some amount of separation, as opposed to pattern convergence^6^ which one could have expected from their reputation of having a much more reliable and precise spiking behavior than principal neurons^40,43^. The high fidelity in relaying input spike^40^ might still explain the difference in pattern separation ability between FSs and GCs, although, to our knowledge, they have never been formally compared. We thus first confirmed the idea that FS show much less spike-wise noise than GCs (**Figure S4**). Linear regressions then revealed that spiking probability (SP) is a good predictor of both spiketrain unreliability (1 – R_w_) and decorrelation performance in FSs (**Figure S3b** and **Table S2**). Surprisingly, the membrane resistance of FSs was also a good predictor (**Table S1**). Thus, contrarily to GCs, FS pattern separation behavior is strongly and linearly determined by some intrinsic and spike-wise properties, even though it is in principle hazardous to anticipate complex neuronal operations from such low-level characteristics, as our previous analysis on GCs illustrated. Indeed, spike-wise noise parameters of HMCs were very close to those of GCs (**Figure S4**) and they nonetheless showed striking pattern separation differences.

Across all celltypes, it is clear that higher SP correlates with lower decorrelation levels (**Figure S3d**). This suggests that sparseness can be a mechanism partially supporting temporal pattern separation. Only partially, because: 1) the correlation is not perfect (R^2^ = 57%), 2) although the correlation is better in FSs, FSs decorrelation levels are consistently lower than expected by the linear model fitting all celltypes (**Figure S3d**), 3) There is no difference in terms of sparseness between CA3 PCs and their GC controls (**Figure S7**) even though there are clear differences in terms of pattern separation (**Figure 4-5**).

In the end, the best predictor of decorrelation levels for all celltypes and conditions is R_w_, the spiketrain reliability (**Figure 8a** and **S6**). This result emphasizes the unexpected idea that the biophysics of neurotransmission in hippocampal networks imposes a trade-off between temporal pattern separation and reliable information transmission. Figure 9 suggests that the balance between separation and reliability can depend on synaptic dynamics. Because different celltypes have synapses with unique properties, we hypothesize that those differences produce varying degrees of pattern separation behaviors not explained by spiking sparseness (SP, FR, pBurst). For example, in contrast to PP-GCs synapses that mostly exhibit depression, the monosynaptic connection from GCs to CA3 PCs is made through giant mossy fiber buttons with low initial probability of release and short-term facilitation under high input frequency^31,44^ (**Figure S9a**). By modelling a spiking neuron with synapses inspired by GC-CA3 connections and comparing it to the model from Figure 9, under conditions yielding similar FR, we confirm that differences in short-term synaptic dynamics alone can lead to obvious pattern separation differences (**Figure S9b-c**).

Overall, these results show that the differences in temporal pattern separation between different hippocampal celltypes result from a combination of various sources, each celltype with a unique combination.

## Discussion

We report that similar cortical input spiketrains are transformed in the DG network, leading to less similar output spiketrains in GCs. Our findings provide the first experimental demonstration that a form of pattern separation is performed within the DG itself and exhibited at the level of single neurons at different timescales. This computation arises from noisy but specific biophysical processes (e.g. synaptic dynamics) in the DG network, where interneurons do not exhibit as much temporal pattern separation as the final DG output. In turn, the CA3 network seems to amplify this separation even more at the level of single PCs, suggesting that, at least in the hippocampus, it is not a computation specific to the DG.

### A novel way to test pattern separation

In contrast to *in vivo* experiments that have difficulty identifying the cell-type of recorded units with certainty^26,45-47^ and simultaneously recording the direct inputs of these units^13-17^, *in vitro* brain slices that preserve the lamellar connections of the hippocampus offer a more accessible platform. For example, a similar experimental setup to ours was used to show that spatially segregated axonal inputs are represented by distinct spatiotemporal patterns in populations of DG neurons^48,49^. However, our study is the first to perform an experimental analysis of pattern separation within DG by directly manipulating the similarity of the inputs and comparing it to the similarity of simultaneously recorded outputs. Such a systematic approach had so far only been done in computational studies^50^. Although a rigorous comparison is impossible because the activity patterns considered were defined differently, the general pattern separation behavior of those models is confirmed by our experimental results: the DG itself performs a form of pattern separation, especially for input patterns that are highly similar (**Figure 2c-d**).

### Pattern separation in the time dimension

Until now, most studies of pattern separation in the DG assumed that neural activity patterns were ensembles of ON/OFF neurons^4,5,15,16,20^, sometimes considering a rate code averaged over minutes in addition to this population code^13,14,23,39^. Because neurons carry information at timescales shorter than minutes^27-29,38,51^ and because the sparse firing of active GCs during a brief event^17,52,53^ precludes an efficient rate code^38^, we studied pattern separation at sub-second timescales.

Relevant scales are given by the time constant over which neurons can integrate synaptic inputs^28^: 10-50 ms for GCs and ∼100 ms for the “reader” CA3 pyramidal cells. Windows of ∼10 ms and ∼100 ms, corresponding respectively to gamma and theta rhythms, have been shown to organize CA neuronal assemblies^27,28,54,55^. Due to specific network properties allowing persistent activity, the DG might also integrate information over several hundreds of milliseconds^48,49,56^. The point is that multiple timescales can be relevant simultaneously, and because it is still uncertain which ones are the most important to episodic memory and hippocampal coding, we investigated a range from 5 ms to 250 ms (**Figure 5**).

Most of our results are reported at a 10 ms resolution, which corresponds approximately to the spike jitter in GCs (**Figure S2**) as well as their membrane time constant and the gamma rhythm. This choice of temporal resolution is similar to a recent computational study of pattern separation within a DG model, which used a 20 ms resolution on short spiketrains (30 ms inputs, 200 ms outputs)^21^. Yet, our study was the first to investigate pattern separation at a range of timescales. We found that temporal pattern separation in the DG output was best at short timescales. This relationship was generally conserved across celltypes, with HMCs even achieving the opposite of pattern separation, pattern convergence, at timescales above 100 ms. Note, however, that temporal pattern separation is not necessarily a monotonically decreasing function of the time resolution, as CA3 PCs exhibited a surprising sharp increase of temporal pattern separation for low input similarity at 250 ms (**Figure 5**).

Our study of pattern separation is the first to focus on temporal patterns, as opposed to spatial ones; but neural activity patterns are spatiotemporal. More work is clearly needed to test whether the DG is a pattern separator at the spatiotemporal, population level, but the discovery of temporal pattern separation in single hippocampal neurons has some implications for population dynamics. The decorrelation of spiketrains at small timescales, in addition to the fact that different GCs respond differently to the same inputs (**Figure S1**), suggests that spikes are constantly rearranged in different time windows, thus enforcing very small neuronal assemblies^28^. In other words, it ensures that a minimal number of output neurons are active at the same time, and such sparsity in active neuronal population is known from computational studies to be critical for efficient population pattern separation^5,22,57^.

### Mechanisms of temporal pattern separation

The mechanisms supporting pattern separation within DG had so far never been experimentally investigated. The decorrelation of sequentially presented input patterns can in theory be explained by: 1) adaptive mechanisms, involving learning and recognition of input patterns, comparison with previously stored ones and the pruning out of common features, 2) non-adaptive (intrinsic) mechanisms, 3) or both ^58^. First, concerning adaptive mechanisms, it has been suggested that Hebbian learning could enhance population pattern separation in the DG^59^, but computational models testing different forms of long-term synaptic learning found that it would actually impair this type of pattern separation^5,23^. As for temporal pattern separation, our data show that it hardly benefits from the repetition of input patterns (**Figure 6**). We also offer indirect evidence that non-adaptive decorrelation processes support temporal pattern separation because output patterns are always decorrelated to the same proportion (**Figure 2e**), a feat that a simple random process can achieve (**Figure S5c**), suggesting that input patterns do not need to be recognized. Third, adaptive and non-adaptive mechanisms are not mutually exclusive: previous learning over days, during the neuron maturation process, could tune single GCs only to specific input patterns, allowing rapid pattern separation^60^. Indeed, a computational study suggested that adaptive networks can mature to perform a fast, non-adaptive orthogonalization of the population activity by the decorrelation between individual information channels^61^.

Adaptive or not, what is the biological source of the temporal decorrelation we observed? We first determined that intrinsic membrane properties do not predict decorrelation levels (**Table S1**), and that celltypes ability to fire bursts only moderately affects pattern separation (**Figure S8**). Simple randomness was not sufficient to reproduce our results, even though the spiking probability, a form of spike-wise neural noise, plays a partial role (**Figure 8** and **S3**). In the end, temporal pattern separation seems most likely supported by specific short-term synaptic dynamics, thanks to the synergy between the probabilistic nature of neurotransmission and its dependency on the spike-timing history imposed by depression or facilitation (**Figure 9, S9**). Various levels of pattern separation/convergence might thus be achieved in different celltypes due to a unique combination of synaptic properties.

Indeed, GCs are embedded in a network of synapses, notably receiving feedforward excitation from the PP, feedforward and feedback inhibition from FSs^43,62^ and both excitation and disynaptic inhibition from HMCs^63^. Thus, although both FSs and HMCs showed some temporal pattern separation (mostly at short timescales), our results suggest that the final DG output is inherited from synaptic interactions between all DG celltypes, resulting in maximal separation at the level of GCs.

More work is needed to clarify the exact role of FSs and HMCs in DG computations. Computational and experimental studies have suggested that HMCs are involved in some forms of pattern separation^17^, ^20^, ^47^. Our finding of pattern convergence in HMCs for dissimilar inputs may appear in contradiction with the recent reports that HMCs spatial representations remap in dissimilar environments^17,26,46,47^, but 1) activity patterns were defined differently and averaged over longer timescales (minutes) and 2) as argued above, remapping is not a direct measure of pattern separation without knowledge of the input patterns. Overall, we show that HMCs can exhibit both separation or convergence, depending on the time resolution and the amount of similarity between input patterns (**Figure 5**). The impact of such a behavior on pattern separation at the level of GCs remains to be studied. Concerning FSs, they exhibit a poor ability to separate spiketrains (**Figure 3, 5**), but the somatic inhibition they provide could be the source of low spiking probability in GCs and thus improve pattern separation by making GCs responses sparse^5,64^. On the other hand, their ability to relay information reliably^43^ (**Figure S4, S6**) and to precisely control spike timing in target neurons^43^ might actually provide a mechanism that counteracts noisiness in GCs, helping them balance effective separation with fidelity of information transmission to CA3.

### The role of sweep-to-sweep variability

Because the brain needs to be able to recognize when situations are exactly the same, our finding that pattern separation occurs even when the same input pattern is repeated (**Figure 7a**) might seem counter-intuitive at first. However, in theory, the separation and the recognition functions do not have to be supported by the same network. The Hebb-Marr framework actually hypothesizes that the CA3 recurrent, auto-associative network is able to recall the original pattern from a noisy input from DG. Even though most computational models that tested the effect of repetition were consistent with the intuitive view^5,21^, this was likely because they used deterministic neurons. A model considering variability across GCs and a probabilistic spiking behavior resulted, as in our experiments, in separation of repetitions of the same pattern^39^.

In the cortex, the well-known variability of single neuron activity between trials is often supposed to be “averaged out” at the population level so that the output of the population is reliable^36^. It is thus conceivable that considering an ensemble of GCs would increase the signal-to-noise ratio. In fact, when we average out the sweep-to-sweep variability, GCs exhibit pattern separation for highly similar patterns but almost no separation for identical ones (**Figure 8d**).

However, this variability, or “noise”, is not necessarily meaningless^36^. Our results suggest it might be a mechanism amplifying pattern separation (**Figure 8**). The variability might even be just apparent, if we consider that when the same input is repeated it is at different points in time: each repetition could be considered as a different event that needs to be encoded slightly differently. The DG would thus meaningfully add some noise to transform input spiketrains so that cortical information about an event is stored in the hippocampus with a unique random time-stamp, consistent with the index theory of episodic memory^65^.

### Pattern separation and pattern completion in CA3

In the Hebb-Marr framework, CA3 is thought to be a recurrent, autoassociative network that can perform pattern completion, and thus can support the recall of a full memory from a partial cue of the original event^4^. Although CA3 recurrent connections are sparser than previously thought^66^, computational models generally confirmed that CA3 could perform pattern completion of population patterns^4,23,66^. Direct experimental evidence are scarce and unclear, both in vitro^67^, and in vivo^32,68,69^. In addition, the fact that neuronal representations of similar environments are more correlated in CA3 than in the DG was thought to constitute indirect evidence^13,14^, but recent reports suggesting that the DG representations previously measured were not coming solely from DG output cells have clouded these initial conclusions^26,46^.

Assuming that pattern completion is a process realized by CA3, this implies that, when presented with different but similar partial cues of the same initial memory, the final output of CA3 should converge towards the same representation. Our finding that CA3 PCs exhibit high levels of temporal pattern separation might then come as a surprise (**Figure 4-5**). Several lines of reasoning could explain this result. First, we focused on temporal patterns in single cells, and it is possible that a network can perform population pattern completion in addition to temporal pattern separation. Actually, different environments can be represented in CA3 by different populations of PCs, or by the same PCs with remapped firing rates^26,46^, which has led some to conclude that CA3 could perform pattern separation^13^. A recent report even showed that single GCs remap much less than CA3 PCs^26^: assuming GCs and PCs receive inputs with similar overlap, this is consistent with our finding that PCs are better than GCs at temporal pattern separation. Second, we tested CA3 under partial block of inhibition in order to allow PCs to fire. Given that the number of active CA3 PCs is generally higher than DG GCs in vivo^5,24,26,46^, it suggests that our experiment may not have modelled physiological conditions well. Third, CA3 is known to be physiologically and functionally heterogeneous along its proximodistal axis^32^, with studies suggesting that the PCs closest to DG perform population pattern separation^68,70,71^. We recorded from PCs at the CA3b/c border, and it is possible that more distal PCs would exhibit less temporal pattern separation. Last but not least, it is important to note that *pattern completion* and *pattern separation* are not opposite, mutually exclusive computations. Pattern separation and its actual opposite, pattern convergence, describe the similarity of multiple patterns from an input network (e.g. EC) compared to the similarity of the corresponding patterns of another network (e.g. DG)^6^. On the other hand, pattern completion is the process, happening over time, of retrieval of a previously learned full pattern in a network (e.g. CA3) from a partial seed pattern in the same network^5^. Our experiments did not test for pattern completion, as the different trains of an input set were not degraded versions of a previously learned pattern. Moreover, pattern separation and completion are complementary in the sense that pattern completion would benefit from initial input patterns being as separated as possible^4,5^. It thus makes sense that CA3 would start by amplifying the separation of input patterns, as is the case in our data, either before encoding or before proceeding to completion of the seed pattern. In fact, our results suggest that CA3 might complement the separation effectuated in the DG, as CA3 is able to perform high decorrelation levels at long timescales (> 100 ms) when the DG doesn’t (**Figure 5**). CA3 could thus ensure that seed input patterns are well separated at all timescales.

## Materials and methods

### Animals and dissection

All experiments were performed in accordance with the National Institute of Health guidelines outlined in the National Research Council Guide for the Care and Use of Laboratory Animal (2011) and regularly monitored and approved by the University of Wisconsin Institutional Animal Care and Use Committee.

Horizontal slices of the ventral and intermediate hippocampus (400 μm) were prepared from the brains of young (p15-25) or adult (p121 ± 15 days) C57BL/6 male mice (Harlan/Envigo). Adult animals were only used for **Figure 9**. Mice were anesthetized with isoflurane, decapitated, and the brain was removed quickly and placed in ice-cold cutting sucrose-based solution (two different compositions were used. For HMC, CA3 pyramidal cells, their GC controls and GCs from adult mice, we used version #2 ^72^). The cutting solutions contained (version #1 / version #2, in mM): 83 / 80 NaCl, 26 / 24 NaHCO_3_, 2.5 / 2.5 KCl, 1 / 1.25 NaH2PO_4_, 0.5 / 0.5 CaCl_2_, 3.3 / 4 MgCl2, 22 / 25 D-Glucose, 72 / 75 Sucrose, 0 / 1 Na-L-Ascorbate, 0 / 3 Na-Pyruvate, bubbled with 95% O_2_ and 5% CO_2_. During the dissection, brain hemispheres were prepared following the “magic cut” procedure ^73^ with α-angle around 10 to 15° and β-angle around 5 to 10° for GC and FS recordings, and with α close to 0° and β between 0 and −5° for HMC and CA3 pyramidal cells recordings ^66^. Slices were cut using a vibratome (Leica VT1000S) then placed in an incubation chamber in standard artificial cerebrospinal fluid (aCSF) containing (in mM) 125 NaCl, 25 NaHCO3, 2.5 KCl, 1.25 NaH2PO4, 2 CaCl2, 1 MgCl2, and 25 D-Glucose (or in a 50/50 mix of standard aCSF and cutting solution. 100% cutting solution for HMC, CA3 recordings and their GC controls, and GCs from adult mice) at 35°C, for 15-30 minutes after dissection. Slices were stored in the incubation chamber at room temperature for at least 30 minutes before being used for recordings.

### Electrophysiology

All recordings were done in standard aCSF adjusted to 325 mOsm, at physiological temperature (33-35 °C). Whole cell patch-clamp recordings were made using an upright microscope (Axioskop FS2, Zeiss, Oberkochen, Germany) with infra-red differential interference contrast optics. Patch pipettes pulled from thin-walled borosilicate glass (World Precision Instruments, Sarasota, FL) had a resistance of 2-5 MΩ when filled with intracellular solution containing (in mM) 140 K-gluconate, 10 EGTA, 10 HEPES, 20 phosphocreatine, 2 Mg_2_ATP, 0.3 NaGTP. For the dataset from adult mice, we used a slightly different recipe: 135 K-gluconate, 5 KCl, 0.1 EGTA, 10 HEPES, 20 Na-Phosphocreatine, 2 Mg_2_-ATP, 0.3 Na-GTP, 0.25 CaCl_2_. Intracellular solutions were adjusted to pH 7.3 and 310 mOsm with KOH and H_2_O. Both recipes yielded similar electrophysiological behaviors.

Recordings were done using one or two Axopatch 200B amplifiers (Axon Instruments, Foster City, CA), filtered at 5 kHz using a 4-pole Bessel filter and digitized at 10 kHz using a Digidata 1320A analog-digital interface (Axon Instruments). Data were acquired to a Macintosh G4 (Apple Computer, Cupertino, CA) using Axograph X v1.0.7 (AxographX.com). Stimulation pipettes were pulled from double barrel borosillicate theta-glass (∼10 μm tip diameter, Harvard Apparatus, Edenbridge, U.K.) and filled with ACSF or a 1M NaCl solution and connected to a constant current stimulus isolator used to generate 0.1-10 mA pulses, 100 microseconds in duration.

Series resistance and cellular intrinsic properties were assessed online in Axograph from the fit of the electrical responses to repetitions of 5-10 mV, 25 ms steps, holding the potential at −65 mV. Neurons used for analysis were stable across a whole recording session as judged by monitoring of series resistance and resting potential.

Dentate granule cells (GC) were visually identified as small cells in the granule cell layer (GCL). GCs from young mice had the following intrinsic properties (mean ± sem) : resting potential (V_rest_) −69.3 ± 1.3 mV; input resistance (R_i_) 171 ± 16MΩ and capacitance (C_m_) 23 ± 2 pF. GCs from adult mice had the following intrinsic properties: V_rest_ −78.8 ± 1.8 mV; R_i_ = 137 ± 14 MΩ and C_m_ = 18 ± 1 pF.

Fast-spiking interneurons (FS) were identified as neurons with large somata at the hilus-GCL border and a high firing rate response during large depolarizing current steps, and a large after-hyperpolarization (AHP) ^62,74^ (**Figure 3**). They had the following intrinsic properties: V_rest_= −66.7 ± 3.5 mV; R_i_ = 59 ± 10 MΩ and C_m_ = 19 ± 3 pF.

Hilar Mossy Cells (HMC) were identified as large neurons in the deep hilus (> 60 µm away from the GCL) with a regular firing response to current steps and a small AHP, as well as with a high frequency of spontaneous EPSPs, all characteristics that allow to distinguish them from other neurons in the hilus ^63,75^ (**Figure 3**). Their intrinsic properties were: V_rest_ = −69.7 ± 2.3 mV; R_i_ = 198 ± 12 MΩ and C_m_ = 33 ± 3 pF.

For CA3 pyramidal cells, because recent studies suggest a proximodistal gradient of physiological and computational properties in the CA3 network ^32,68,70,71^, we avoided the extremity of the CA3c region and targeted large neurons around the CA3b/c border in the pyramidal layer. Their intrinsic properties under partial inhibitory block (aCSF + 100 nM gabazine) were: V_rest_ = −72.7 ± 2.2 mV; R_i_ = 186 ± 12 MΩ and C_m_ = 36 ± 2 pF. The intrinsic properties of GCs recorded under the same pharmacological conditions were: V_rest_ = −77.3 ± 1.9 mV; R_i_ = 244 ± 13 MΩ and C_m_ = 25 ± 2 pF.

### Pattern separation experiments

We designed multiple sets of input patterns, each with a prespecified average Pearson’s correlation coefficient (R_input_) computed with a binning window (τ_w_) of 10 ms, using two different algorithms (one developed in-house based on iterative modifications, and a more efficient one from ^76^ based on mathematical principles and a preset covariance matrix). An input set consisted of five different input spiketrains, each 2 s trains of impulses simulating cortical spiketrains, with interspike intervals following a Poisson distribution. Each pair of input trains had a correlation coefficient close to the R_input_ of its set (at τ_w_ = 10 ms, the average relative standard error is 4% across all input sets). For all experiments except in **Figure 4** (CA3 and GC control), the mean frequency of input trains was set close to 10 Hz (11.9 ± 0.7 Hz). This input firing rate was chosen to be consistent with the frequency of EPSCs recorded in GCs of behaving mice ^52^, and is known to promote a high probability of spiking in GCs in slices ^62,77^.

The responses of one or two DG neurons were recorded in whole-cell mode while stimuli were delivered to the outer molecular layer (OML). Stimulus current intensity and location were set so that the recorded neuron spiked occasionally in response to electrical impulses (see range of spike probability in **Figure S2-4**) and the stimulation electrode was at least 100 µm away from the expected location of the dendrites of the recorded neuron. Once stimulation parameters were set, a pattern separation protocol was run: the five trains of a given input set were delivered one after the other, separated by 5 s of relaxation, and this was repeated ten times. The ten repetitions of the sequence of five patterns were implemented to take into account any potential variability in the output, and the non-random sequential scheme was used to avoid repeating the same input spiketrain close in time. Each protocol yielded a recording set of fifty output spiketrains, each associated with one of the five input trains of an input set (**Figure 1c**). A given cell was recorded in response to up to five input sets with different R_input_ (i.e. a recorded cell produced between one and five data points on **Figure 2c**).

The membrane potential baseline was maintained around −70mV during both current-clamp and voltage-clamp recordings, consistent with the V_rest_ of mature GCs recorded in behaving mice ^52^. For comparison, FS and HMC current-clamp recordings were also held at −70 mV. The output spiking frequency of GCs was variable (6.3 ± 0.3Hz, see **Figure S7-8**) but consistent with sparse activity generally observed in GCs *in vivo* during behavior ^26,52,53,78^ and in slices under conditions of drive comparable to ours ^41,79,80^.The output firing rates of FS and HMC were higher than their GC controls (**Figure S7-8**), as expected from recent research ^26,43,47^.

To perform pattern separation experiments in CA3 pyramidal cells (PCs), we had to change several parameters in order to make CA3 PCs spike. Indeed, PCs firing is controlled by strong feedforward inhibition ^31,32^, and all tested stimulation sites led to net IPSPs or, rarely, to weak EPSPs. To make PCs fire in response to external electrical stimulations: 1) the stimulating electrode was placed in the inferior blade of the GCL to make the DG output cells fire, 2) we targeted CA3b PCs, which have the highest E/I ratio across CA3 in slices and still receive strong connections from mossy fibers ^32^, 3) we used input sets with 30 Hz input trains to promote the depression of inhibitory transmission ^31^, 4) we maintained the membrane potential baseline between −60 and −70 mV, and 5) we had to add 100 nM gabazine to the bath of standard aCSF in order to slightly decrease IPSCs amplitude (recordings started at least 15 min after the slice was placed in the bath, to insure equilibrium was reached). 100 nM of gabazine (SR-95531, Sigma-Aldrich) has been shown to correspond to a 70% availability of GABA-A receptors ^81^ and, in our conditions, consistently decreased spontaneous IPSCs mean amplitude by 30% in voltage-clamp recordings of GCs and CA3 PCs held at −40 mV. These conditions allowed us to record for the first time the spiking output of CA3 PCs in response to complex input spiketrains while preserving some inhibition in the network ^31^. PCs output firing rate were on average below 10Hz (**Figure S7-8**), but close to mean rates observed in vivo during behavior ^13,14,26^.

### Analysis of the output spiketrains

For each recording set, the similarity between pairs of spiketrains was computed as the Pearson’s correlation coefficient between the spiketrains rasters binned at a τ_w_ timescale. Sweeps without spikes were excluded from further analysis.

We did not use separate protocols to assess the *firing rates, probability of bursting* and *spike-wise noise* parameters (spike probability, delay, jitter), but computed them directly for each recording set of spiketrains from a pattern separation experiment. The mean firing rate was computed as the average firing rate across all fifty output spiketrains. A burst was defined as the occurrence of more than one output spike in the interval of two input spikes (see **Figure S8**).

We define the spike-wise neural noise as the *probability of spiking* at least once after an input spike, the *delay* of an output spike after an input spike and its average *jitter*. To assess these parameters, we computed the cross-occurrence between input spikes and output spikes in a [-15 ms, 50ms] interval with 1 ms bins. The resulting histogram of counts of output spikes occurring in the vicinity of an input spike was fitted with a Gaussian distribution *N*(µ,σ, baseline), where µ is the mean delay of an output spike and σ is the jitter of this delay. The baseline corresponds to the background firing, occurring by chance or caused by neighboring inputs. After subtracting the baseline and extracting the probability of spiking by dividing the counts of output spikes by the total number of input spikes, we defined the *spike probability* (SP) as the sum of probabilities of an output spike in the predefined time interval around an input spike (**Figure 7b** and **Figure S2a**). For FS and HMC, spike-wise noise parameters were computed on a non-bursty dataset (nbFS and nbHMC) where all but the first spike in a burst were excluded. Spike-wise noise parameters were not computed for CA3 PCs and their GC control because the input frequency was too high to allow a good fit of the distribution.

### Computational models

To assess the role of spike-wise neural noise in pattern separation, we generated two data sets. First, we simulated output spiketrains in response to our input sets (for each input set, we simulated ten output sets of fifty synthetic spiketrains). This simulation was entirely based on the average spike-wise noise parameters computed from the original GC recordings (see above): the matrix of input spike times was replicated ten times, and for each of the fifty resulting sweeps, spikes were deleted randomly following a binomial distribution *B*(Nspk, F), where Nspk is the number of input spikes in a sweep, and F the probability of not spiking (F = 1-mean SP = 1-0.42). A random delay, sampled from a Gaussian distribution *N*(µ,σ), was added to each resulting spike times, with µ and σ being respectively the mean delay and mean jitter in the original recordings. The noise statistics of the resulting simulated data set is shown in **Figure S2c**. Second, we created a surrogate data set by randomly shuffling the output spikes of the original GC recordings: the delay of each spike was conserved but it was relocated to follow a randomly selected input spike in the same input train (from a uniform distribution). This manipulation yielded a dataset with noise statistics very close to the original data (**Figure 2d)**. Using this strategy, we performed spike shuffling in each GC recording set a hundred times, yielding a dataset of 10,200 simulated recording sets, or, in other words, 100 datasets of 102 simulated recording sets directly paired to the original 102 GC recording sets.

To test whether probabilistic synaptic dynamics is a potential mechanism of temporal pattern separation, we used an Izhikevich model of a regular spiking neuron ^82^ with an adapted Tsodyks-Markram (TM) synapse model ^83^ designed to capture short-term plasticity of stochastic neurotransmission at the LPP-GC synapse. The TM synapse model consists of a system of two ordinary differential equations describing the dynamics of X and U. X is the probability of a presynaptic vesicle being available for release, the decrease of which leads to synaptic depression, and U is the probability of release for an available vesicule, the increase of which models facilitation:

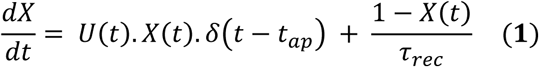

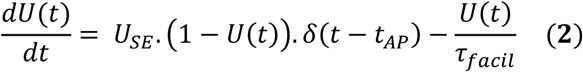

With X(0) = 1 and U(0) = 0, where τ_rec_ is the time constant of vesicule recovery after release, τ_facil_ is the decay of U controlling synaptic facilitation, U_SE_ a factor determining the probability of release Pr at the time of the first spike, δ the Dirac delta function, and t_AP_ the time of arrival of an input spike. The system of ODEs was solved using the ode23 MATLAB solver.

Based on the quantal theory of neurotransmission, the model assumes that, at each time point t, *k* vesicles are released, drawn from a binomial distribution *B*(N, Pr), where N is the number of release sites and Pr is the probability of release defined as the product of X and U. The current *I* is then the product of *k* with the quantal size *q*. Based on previous data from our lab, we considered *q* = 20 pA ^62^. The version of the TM model we implemented only focuses on the dynamics of the presynaptic vesicles, ignoring postsynaptic dynamics. To match the decay of real EPSCs recorded at the soma, *I* was convolved with a template EPSC shape:

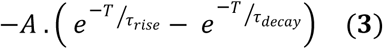

Where T = 50 ms, τ_rise_ = 0.5 ms, τ_decay_ = 5 ms and A is a factor normalizing the area of the resulting shape to 10. In addition, a synaptic delay of 4 ms was introduced. The resulting current (**Figure 9b-c**) is then passed to a non-dimensional regular spiking Izhikevich neuron, an acceptable model for GCs ^84^. We used the following parameters: a = 0.02, b = 0.2, c = −65 mV, d = 6, time resolution = 0.1 ms, max voltage = 30 mV, initial values V(0) = −70 mV and x(0) = b.V(0).

To optimize the non-fixed parameters of the TM-Izhikevich model, we simulated output spiketrains in response to ten repetitions of a single input set (10 Hz, R_input_ = 0.76) and compared to the current dynamics and spiking behavior of a representative GC that was recorded both in voltage-clamp and current-clamp in response to the same input set. For the TM model, the following parameters were adjusted to match the dynamics and the peak of the current averaged over ten repetitions, as well as the variability in the peak amplitude of the first EPSC: τ_rec_ = 500 ms, τ_facil_ = 9 ms, U_SE_ = 13, N = 40 release sites. Finally, because the Izhikevich model integrates non-dimensional currents, the input current had to be scaled using the dividing constant K = 62, such that the average firing rate of the simulation matched the average firing rate of the representative GC to the same input set (3.5 Hz). K can be considered as a constant modelling a tonus of inhibition: the higher K is, the more difficult it is for an EPSC to make the Izhikevich neuron fire. In the end, the standard deviation of the firing rate was ∼0.6 Hz for simulations, compared to 0.8 Hz for the original recording. The only source of variability was from presynaptic dynamics of vesicle release modelled as a binomial process.

To determine the influence of synaptic transmission parameters on pattern separation, we compared two TM-Izhikevich models with different parameters. Model 1 has exactly the same parameters as described above except that, in order to model the presence of gabazine, the inhibition constant K was set at 40: with this value, Model 1 responses to 30 Hz input trains have a FR ∼ 7 Hz, corresponding to the mean FR of real GCs recorded under partial inhibitory block (**Figure S7**). For model 2, q = 29 pA, τ_rec_ = 100 ms, τ_facil_ = 500 ms, U_SE_ = 0.01, K = 57, and all other parameters were the same as for model 1. U_SE_, q and N (kept at 40) were chosen based on estimates from the literature on mossy fiber buttons, the giant GC-CA3 PCs synapses^31,44^. τ_rec_, τ_facil_ and K were adjusted to model short-term synaptic facilitation observed at mossy fiber buttons and to match the average firing rate that we observed in our CA3 current-clamp recordings made under gabazine (∼7Hz, **Figure S7**). In order to specifically study the impact of pre-synaptic dynamics, post-synaptic (i.e. EPSC shape) and Izhikevich parameters were kept the same for all models. Note that none of our models are intended to closely match spiking behaviors observed in real GCs or CA3 PCs.

### Software and statistics

Data analysis was performed using custom-written routines in MATLAB (2017a), including functions from toolboxes cited above. Sample sizes were chosen based on the literature and estimations of the variance and effect size from preliminary data. All values are reported as mean ± S.E.M. unless otherwise noted. The one-sample Kolmogorov-Smirnov test was used to verify the normality of data distributions. Parametric or non-parametric statistical tests were appropriately used to assess significance (p-value < 0.05). Assumptions on equal variances between groups were avoided when necessary. All T and U tests were two-tailed. To determine whether two distributions of data points are significantly different (e.g. R_output_ as a function of R_input_, for GC compared to FS, see **Figure 3-6 and Figure S5 and S8**), we performed an analysis of the covariance (ANCOVA) using linear regressions on the two data sets as well as on the combined data set, and assessed significance via an F-test comparing the goodness of fits^85^. Because R_input_ can also be considered as a categorical variable, we performed a two-way ANOVA before using post-hoc tests correcting for multiple comparisons in order to determine at which R_input_ groups two conditions were significantly different. When comparing FSs, HMCs or CA3 with GCs, different GCs were used as control and were recorded under different protocols, which is why we did not need to control for multiple comparisons across celltypes. In order to determine whether distributions were significantly different in the case of our spike shuffling analysis (**Figure 7**), we designed Monte-Carlo exact hypothesis tests^86^. **Table S4** provides details on all statistical tests conducted in this study.

### Data availability

Data used in this manuscript are freely available at: https://www.ebi.ac.uk/biostudies/studies/S-BSST219

## Supplements

Tables and supplementary figures for “Pattern separation of spiketrains in hippocampal neurons” Authors: A.D. Madar, L.A. Ewell, M.V. Jones.

**Figure S1.**
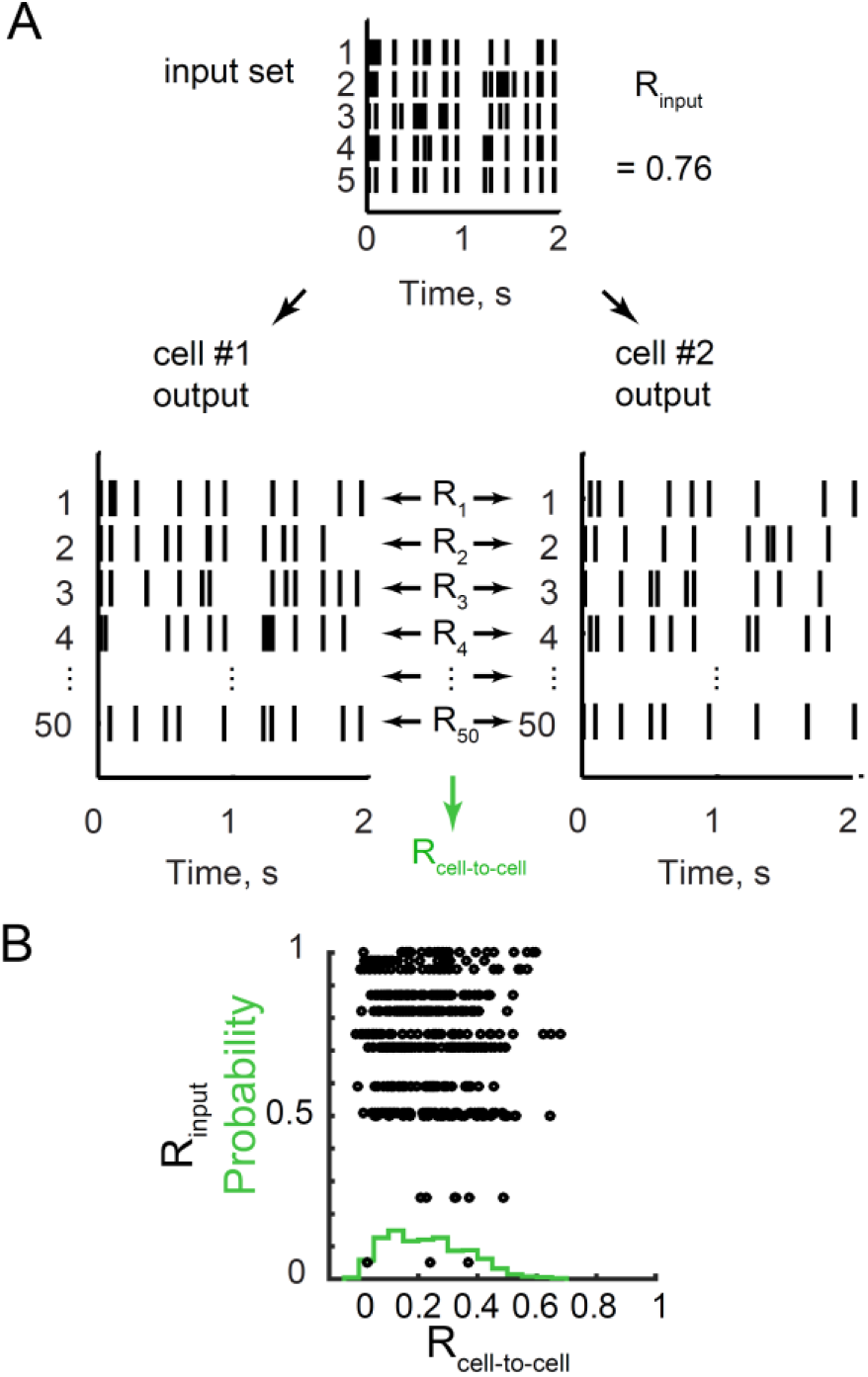
Different granule cells process identical inputs differently. **(A)** The similarity between pairs of spiketrains coming from two different output sets but associated to the same input set and with the same sweep number is assessed with the Pearson’s correlation coefficient (τ_w_= 10 ms). The fifty resulting coefficients are then averaged to give R_cell-to-cell_, a single number measuring the overall similarity of all output spiketrains between two output sets. **(B)** Probability distribution of R_cell-_ to-cell (green histogram) across all GC recordings (all combinations of pairs of GC output sets from the same input set were compared). The distribution of R_cell-to-cell_ (black circles) is not dependent on R_input_.

**Figure S2.**
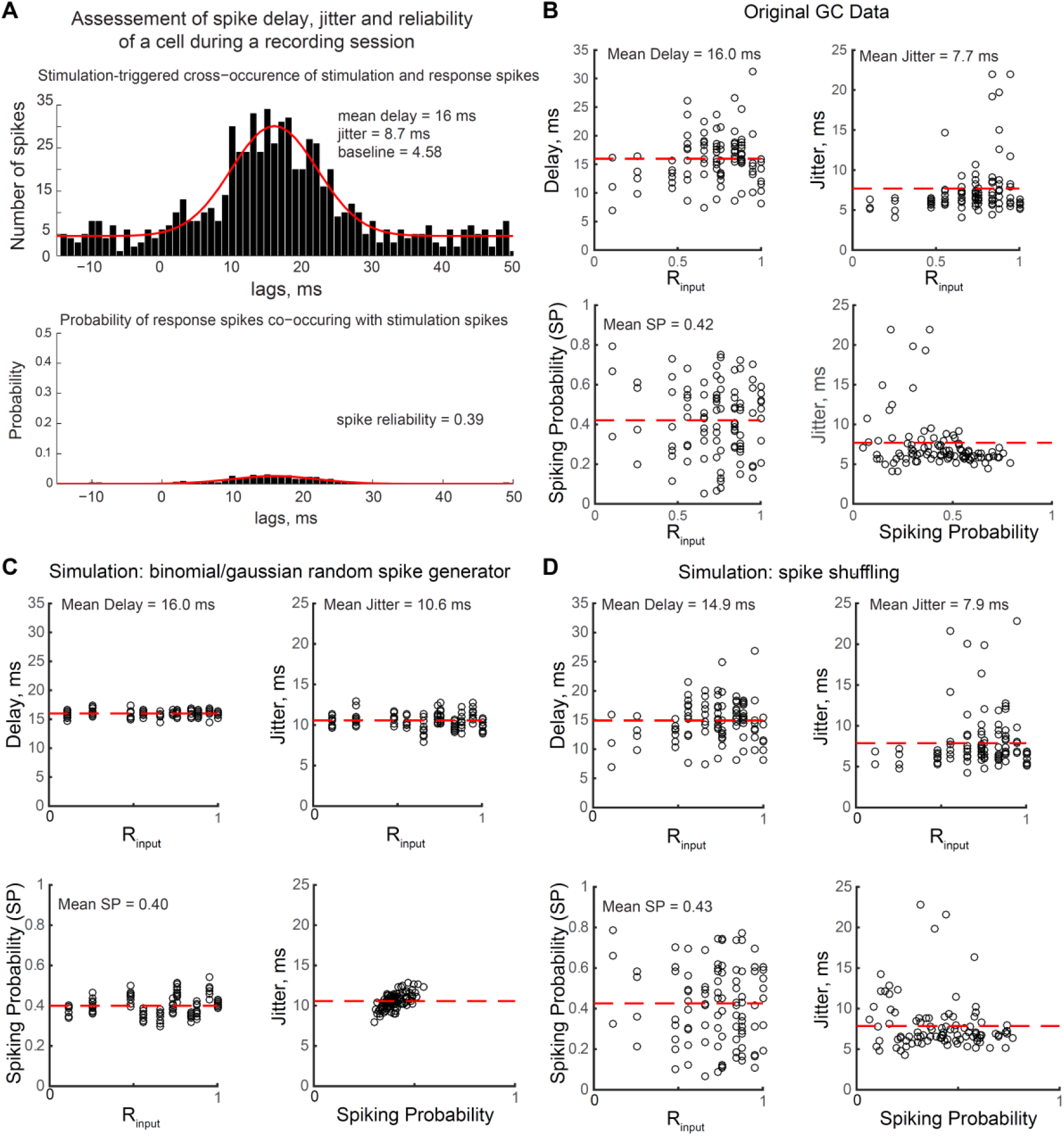
related to Figure 7. Spike delay, jitter and reliability distributions for real data, simulations and shuffled data. **(A)** Cross-occurrence method to measure spike delay, jitter and spiking reliability of a neuron during a given recording session. *Top*: Example histogram of output spikes occurring after input spikes, fitted (red curve) with a Gaussian distribution N(µ,σ, baseline), where µ is the mean delay and σ is the jitter of this delay. Lag 0 ms corresponds to the input spike time. In this example, output spikes are generated on average 16 ms after a stimulation impulse (delay) with a jitter (σ) of 8.7 ms. *Bottom*: the baseline is subtracted and the histogram divided by the number of input spikes during the recording session. This gives the distribution of the probability of spiking after an input spike, the sum of probabilities defining the spiking probability (SP) of the cell during the recording session. Here the neuron fires 39 % of the time after an input spike. **(B-D)** Delay, jitter and spiking probability (SP) distributions as a function of input sets, for (**B**) the original GC recordings, (**C**) the simulations of binomially random spiking with Gaussian delay (n = 110) and (**D**) one spike shuffling dataset (n = 102). Dashed horizontal red lines are means.

**Figure S3.**
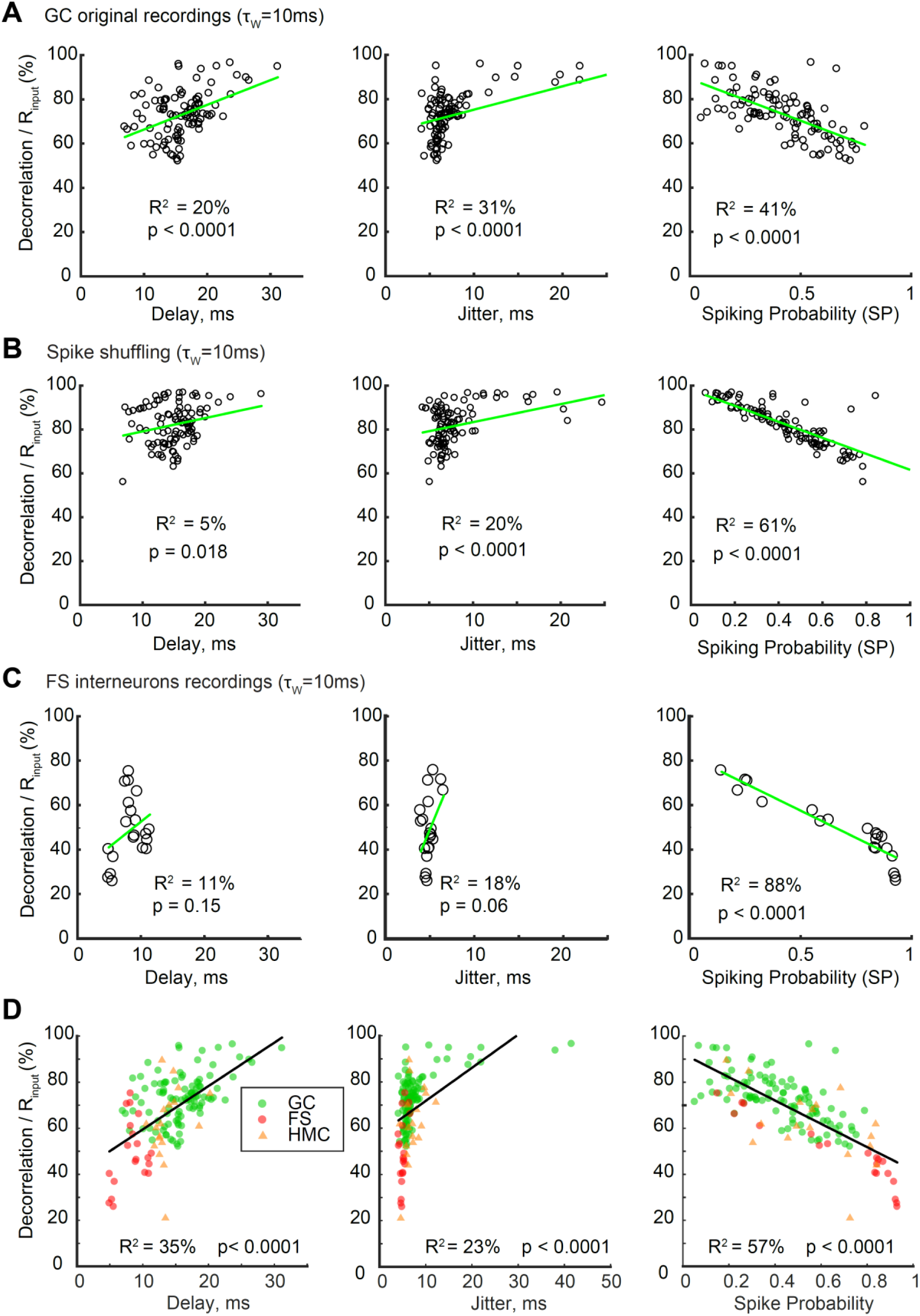
related to Figure 7. Spike-wise neural noise characteristics are not good predictors of spiketrain decorrelation by single GCs. Plots of the normalized decorrelation, i.e. (R_input_ – R_output_) / R_input_, of each recording set (τ_w_ = 10 ms) : 102 for GC original and shuffled recordings **(A-B)**, 20 for FS **(C)**, and for GC, FS and HMC pooled together (**D**) as a function of spike-wise noise characteristics (spike delay, jitter and probability). Solid green lines are the best linear fit, with R^2^ and p-values noted in each panel. These plots illustrate **Table S2**. Note that decorrelation is poorly explained (low R^2^) by either the spike delay or its jitter in all cell-types. In contrast, the spiking probability (SP) is a good predictor of decorrelation in shuffled GC recordings (n = 102 recordings entirely dominated by spike-wise noise. See Figure S2) and even more so in FS recordings (for FS, SP was computed from nbFS data. See Figure S7). This suggests that a low SP can be a potent mechanism for decorrelation, and that FS show different levels of decorrelation than GCs partly because they are more reliable. However, the regression line for FS is lower than for GCs (even FS with low SP show less decorrelation than GCs with similar SP), and SP is only an average predictor of decorrelation for GCs, thus confirming that temporal pattern separation in single GCs cannot be the result of simple neural noise.

**Figure S4.**
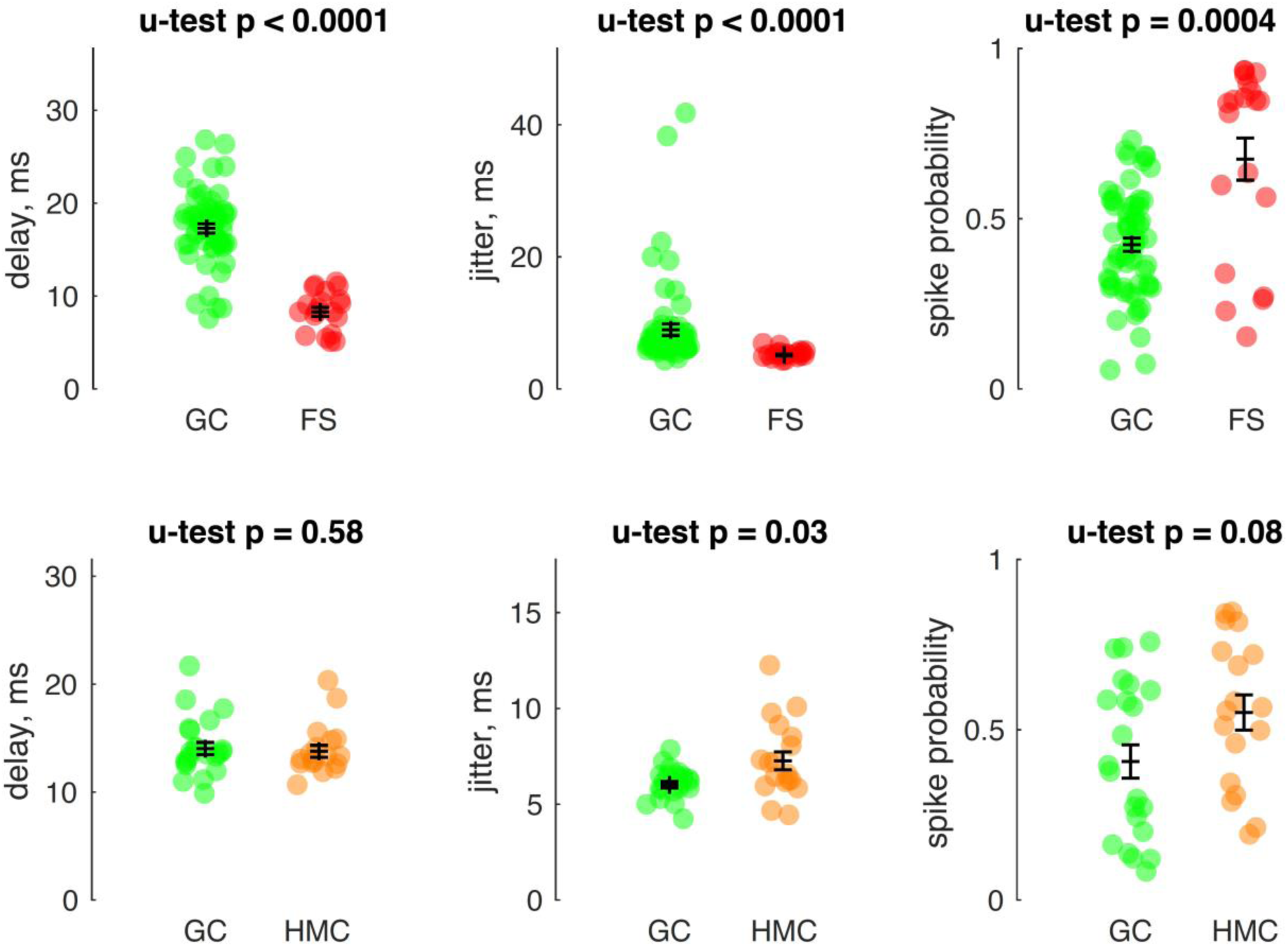
related to Figure 7. Spike-wise noise characteristics of granule cells are different than those of DG interneurons. Mean +/-SEM (bars) and individual recordings (dots). Spike delay, jitter and spiking probability (SP) for FS interneurons compared to GC recordings associated with the same input sets than FS (*top*) (20 FS recordings, 61 GC recordings, see Figure 3A) and HMC interneurons compared to a different set of GC recordings associated with the same input sets than HMC (*bottom*) (18 HMC recordings, 22 GC recordings, see Figure 3B). A U-test was applied to each pair of comparison, showing that FS are much faster and less noisy than GCs, whereas HMC are similar to GCs (slightly larger jitter, but slightly higher SP).

**Figure S5.**
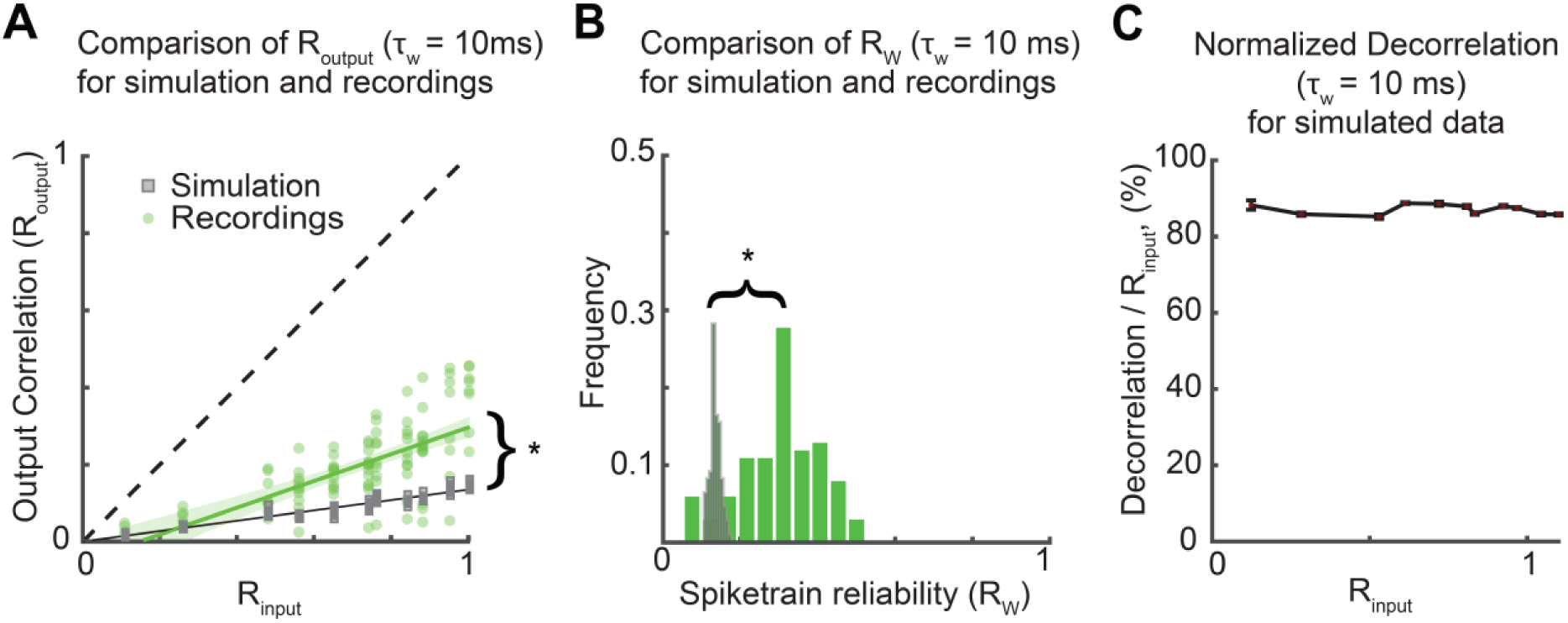
related to Figure 7. R_output_ and spiketrain reliability are lower for a random spike generator than for GCs. A Binomial/Gaussian random spike generator with parameters based on the mean spike-wise neural noise measured in GCs leads to more decorrelation but less spiketrain reliability than in GCs. **(A)** R_output_ distribution at τ_w_ = 10 ms, for simulated and GC recordings. (ANCOVA: p < 0.0001). Shaded areas (green and grey) represent the 95% CI of the regressions. **(B)** R_w_ distributions are significantly different (unpaired T-test correcting for unequal variances: p < 0.0001, <Rw>simul = 0.14). **(C)** Like in the original data (**Fig 2E**), the average normalized decorrelation ((R_input_-R_output_)/R_input_) seems invariant. Bars are SEM. **(A-B)** Asterisks: p < 0.05.

**Figure S6.**
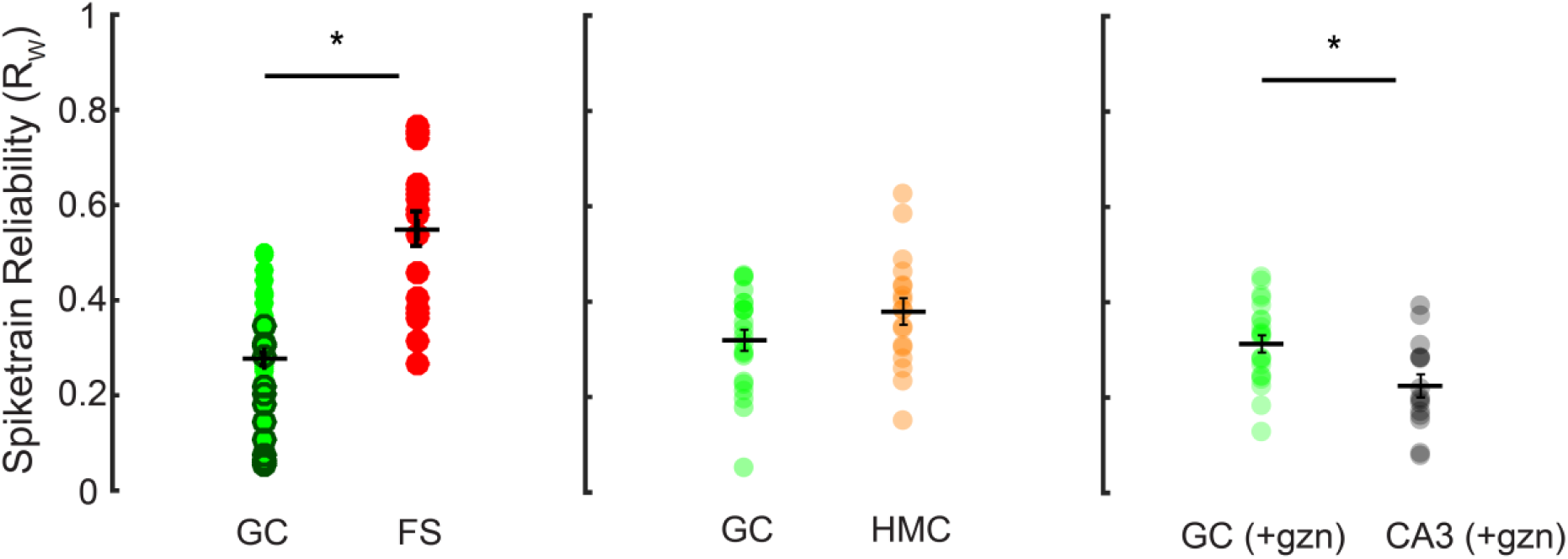
related to Figure 8A. Spiketrain reliability in different hippocampal celltypes. FS have a higher R_w_ than GCs (n = 20 vs 61 recording sets, unpaired T-test: p < 0.0001). Note that when comparing only the simultaneous FS and GC recordings (dark green), we found a similarly significant difference. HMCs R_w_ are not significantly higher than in GCs (n = 18 vs 22 recording sets, unpaired T-test: p = 0.0963). Under partial block of inhibition, CA3 pyramidal cells have a lower R_w_ than GCs (n = 15 vs 22 recording sets, unpaired T-test: p = 0.0052). Asterisks: p < 0.05

**Figure S7.**
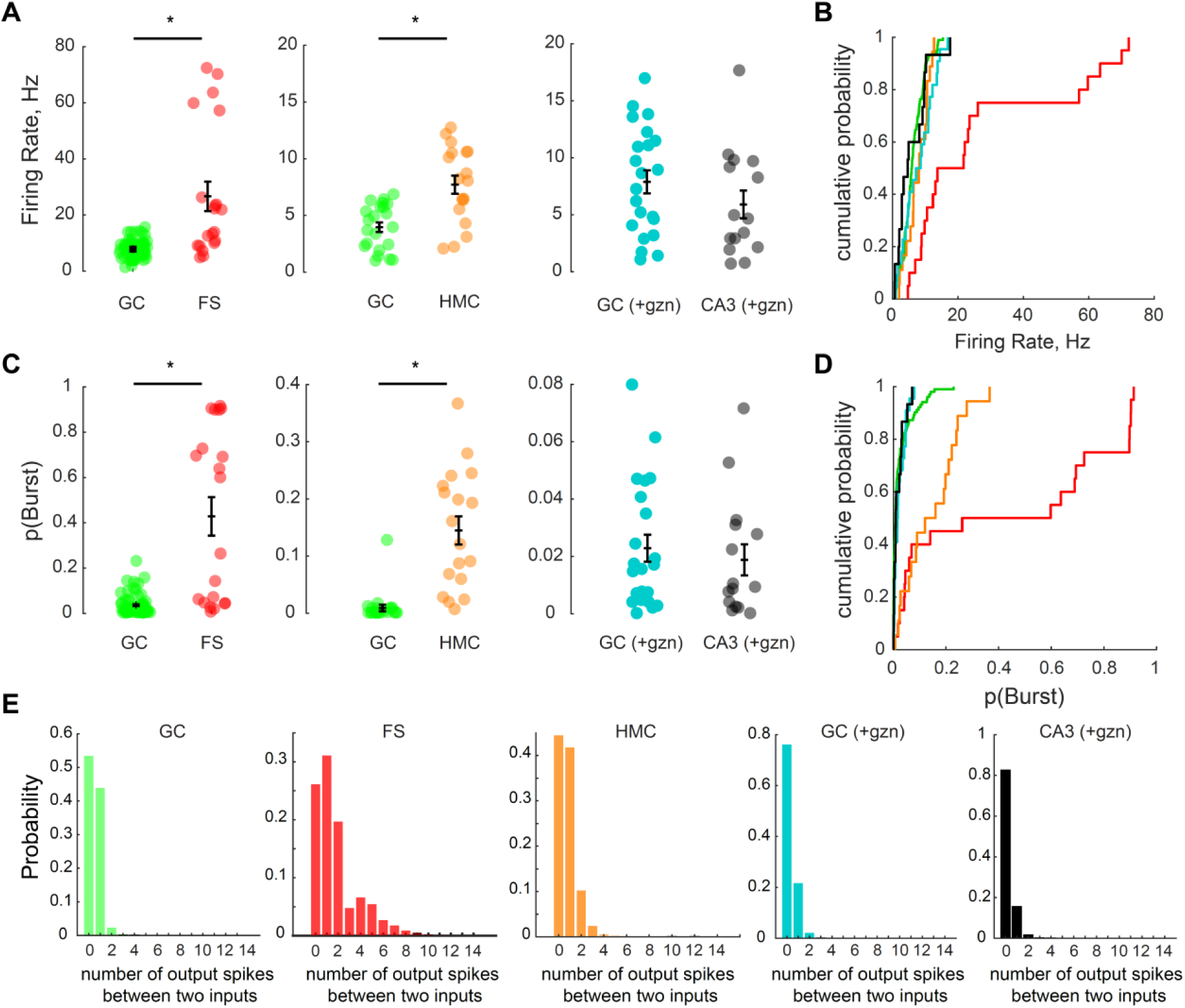
related to Figure 3. DG interneurons fire bursts and have a higher firing rate than GCs. **(A-B)** Comparison of the firing rate in all celltypes and conditions. **(A)** For each comparison the GC dataset is different and matched to the non-GC dataset, as in Figure 3-4. FS and HMC have a higher firing rate than their GC control (U-test: p < 0.0001 and p = 0.0005 respectively) whereas CA3 and GC recordings under gabazine and 30 Hz input trains do not differ (U-test: p = 0.18). **(B)** Cumulative distributions of the firing rate for all recordings of all celltypes (n = 102 GC, 20 FS, 18 HMC, 15 CA3+gzn, 22 GC+gzn). **(C-D)** Same as A-B for the probability of bursting in a recording. Bursting was defined as the occurrence of at least two action potentials between two input pulses. **(C)** FS and HMC have a higher propensity to fire bursts than their GC control (U-test: p < 0.0001) whereas CA3 pyramidal cells and their GC control do not differ (U-test: p = 0.74). **(E)** The distribution of the number of spikes between two input pulses, for all celltypes, show that FS are the only neurons that consistently fire large bursts. Note that GCs almost never fire more than twice, and when they do it generally corresponds to two preceding input pulses occurring close to each other (the first spike being in reality associated to a different input than the next spike).

**Figure S8.**
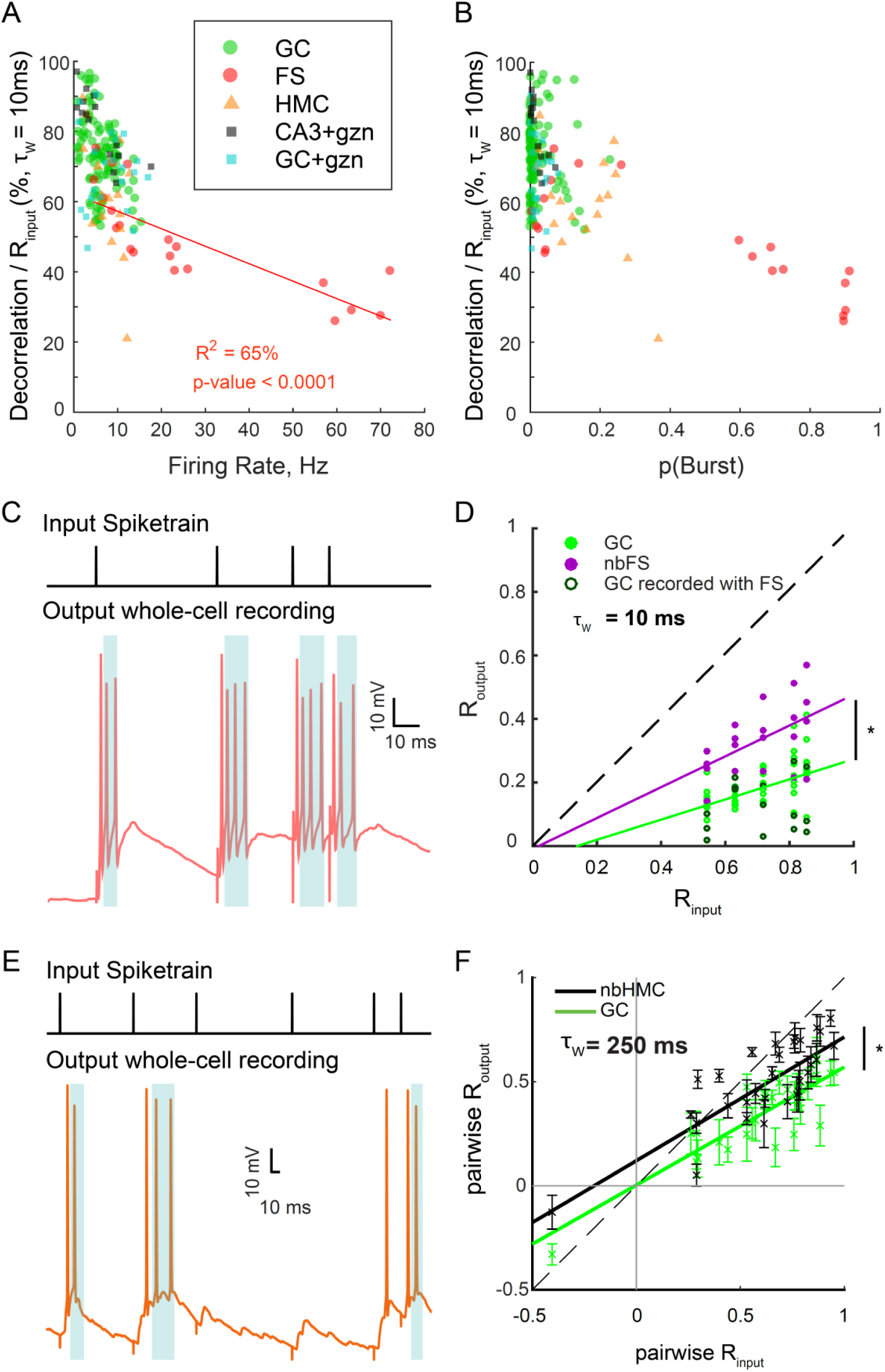
related to Figure 3 and 5. Differences in pattern separation between GC and DG interneurons are not solely due to different firing rates and bursting behaviors. **(A)** The relationship between the firing rate and decorrelation levels in all recordings of all celltypes. In contrast to GCs, for FS neurons there is a strong correlation between the firing rate of a recording set and the associated normalized decorrelation. See **Table S3** for more details. **(B)** The relationship between the probability of bursting (see Figure S7) and the decorrelation levels in all recordings of all celltypes is less clear, but cells bursting more than 30% of the time have the lowest decorrelation levels. **(C)** Example of bursts in a FS recording (*bottom*) in response to input spikes (*top*). To assess the effect of bursting on R_output_, we truncated each recorded spiketrain from FS neurons to keep only the first output spike between two input spikes, thus removing any burst without altering the SP of the cell. The blue shaded areas highlight the spikes that were removed. The resulting truncated dataset was termed “nbFS” for “non-burst FS”. **(D)** R_output_ versus R_input_ for nbFS and GC at τ_w_ = 10ms (to compare to Figure 3A4). Both distributions are still significantly different, suggesting the bursting behavior of FS is not sufficient to explain the difference in pattern separation: ANCOVA: p < 0.0001. **(E)** Bursts in HMC recordings were truncated to produce an “nbHMC” dataset. **(F)** Pairwise analysis on nbHMC and GC recordings at τ_w_ = 250 ms show that the distributions are still different between the two celltypes (to compare to Figure 5D): ANCOVA: p < 0.0001. Under this analysis, nbHMCs and GCs are also significantly different at lower timescales (τ_w_ = 100, 50 and 10 ms) but with a lower effect size as τ_w_ decreases (not shown).

**Figure S9.**
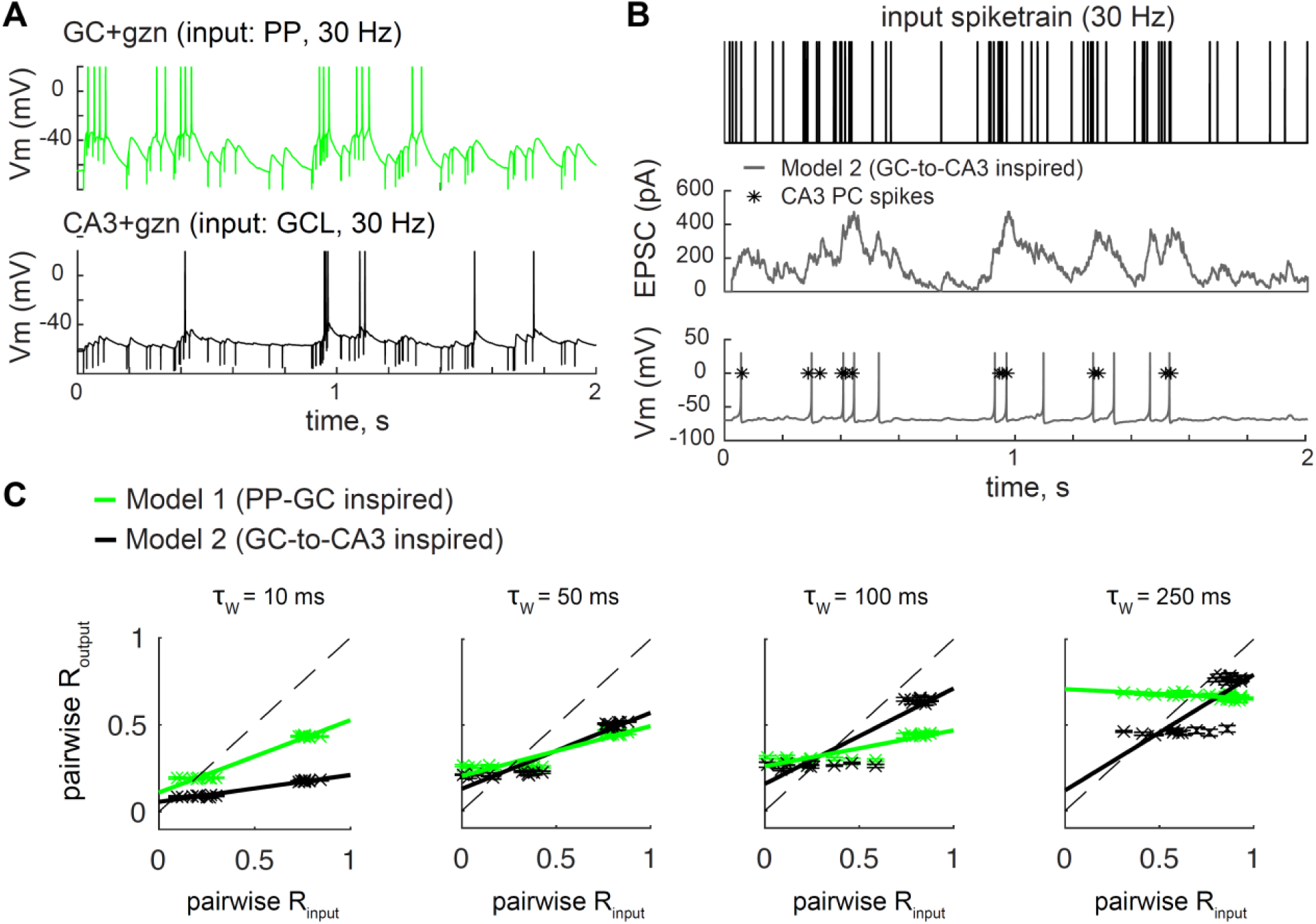
related to Figure 5 and 9. Short-term synaptic dynamics differences can drive differences in terms of temporal pattern separation. **(A)** Examples of current-clamp recordings in one GC and one CA3 PC, under partial block of inhibition, in response to a 30 Hz input train (input pulses are noticeable as downward artefacts). EPSPs visually appear quite different between GCs and CA3 PCs, with clear facilitation in CA3 PCs, which leads them to spike mostly during periods of high input frequencies. **(B-C)** We designed two models of spiking neurons only differing by a few synaptic parameters. Model 1 is the same as the model presented in Figure 9, with depressing EPSC dynamics, except that the inhibition constant was decreased to match FR observed in real GC+gzn recordings. Model 2 was inspired by GC-to-CA3 mossy fiber synapses that exhibit low initial probability of release and short-term facilitation, and the inhibition constant was set so to match the mean FR of real CA3+gzn recordings (∼7Hz). Model 1 and 2 were thus designed to have different synaptic dynamics but yield similar FR. Spiking responses to the pattern separation protocols used on real CA3 PCs and their GC controls (30 Hz input trains: R_input_ = 0.21 and 0.76, shown in Figure 4) were then simulated (n = 5 simulated output sets for each model and each Rinput). **(B)** Example of current and voltage responses of model 2 to a 30 Hz input train. Asterisks correspond to spike times of a single sweep from a CA3 PC recording (different than in A). **(C)** Pairwise correlation analysis, as in Figure 5, shows that the two models yield visually obvious differences in terms of temporal pattern separation (crosses and bars: mean +/-SEM; solid lines: linear regression on data points). Although model 1 and 2 do not reproduce well the pattern separation levels observed in real GCs and CA3 PCs (Figure 5E), they qualitatively go in the same direction: the GC-to-CA3 inspired model 2 produces visually lower R_output_ than the PP-GC inspired model 1, at short timescales (10 ms) and large ones (250 ms, for low R_input_ values). Overall, this analysis shows that differences in terms of synaptic dynamics are sufficient to cause differences in terms of temporal pattern separation.

**Table S1-3**. **Linear regressions goodness-of-fit, p-value and slope.** The predictor variables (x-axis) correspond to columns, and the variables to be explained (y-axis) correspond to rows. Red highlights significant regressions that explain more than 50% of the variance (R^2^ > 50%). Blue highlights regressions that are significant (p < 0.01) but that explain less than 50% of the variance. The values used for Normalized Decorrelation, i.e. (R_input_ - R_output_)/ R_input_, and for Spiketrain Reliability (R_w_) were computed with a binning window of 10 ms, unless specified. Abbreviations: GC yo = from young mice, GC ad = from adult mice, ALL = dataset pooling all celltypes and conditions recorded in young mice

**Table S1.** Intrinsic electrophysiological cell properties

| y-axis ↓<br>x-axis → |  | Membrane Capacitance (Cm) | Membrane Resistance (Rm) | Membrane Time Constant = Rm.Cm | Resting Membrane Potential (Vrest) |
| --- | --- | --- | --- | --- | --- |
| Normalized Decorrelation | GC yo | R<sup>2</sup> = 4%<br>p = 0.08<br>slope = -0.2<br>F(1,100) = 3.2 | R<sup>2</sup> = 5%<br>p = 0.06<br>slope = -0.03<br>F(1,100) = 3.7 | R<sup>2</sup> = 8%<br>p = 0.013<br>slope = -1.2<br>F(1,100) = 6.5 | R<sup>2</sup> = 3%<br>p = 0.17<br>slope = -0.2<br>F(1,100) = 1.9 |
|  | FS | R<sup>2</sup> = 47%<br>p = 0.0008<br>slope = -1.5<br>F(1,18) = 15.8 | R<sup>2</sup> = 77%<br>p < 0.0001<br>slope = 0.6<br>F(1,18) = 58.9 | R<sup>2</sup> = 5%<br>p = 0.4<br>slope = 13.6<br>F(1,18) = 1.2 | R<sup>2</sup> = 46%<br>p = 0.0009<br>slope = 1.4<br>F(1,18) = 15.6 |
|  | HMC | R<sup>2</sup> = 33%<br>p = 0.013<br>slope = -0.4<br>F(1,16) = 7.8 | R<sup>2</sup> = 3%<br>p = 0.48<br>slope = 0.06<br>F(1,16) = 0.5 | R<sup>2</sup> = 7%<br>p = 0.3<br>slope = -1.8<br>F(1,16) = 1.6 | R<sup>2</sup> = 0.5%<br>p = 0.8<br>slope = 0.1<br>F(1,16) = 0.08 |
|  | CA3 +gzn | R<sup>2</sup> = 0.1%<br>p = 0.9<br>slope = -0.05<br>F(1,13) = 0.01 | R<sup>2</sup> = 2%<br>p = 0.59<br>slope = 0.03<br>F(1,13) = 0.3 | R<sup>2</sup> = 4%<br>p = 0.60<br>slope = 1.2<br>F(1,13) = 0.6 | R<sup>2</sup> = 29%<br>p = 0.04<br>slope = 0.7<br>F(1,13) = 5.4 |
|  | GC +gzn | R<sup>2</sup> = 30%<br>p = 0.008<br>slope = 1.6<br>F(1,20) = 8.6 | R<sup>2</sup> = 0.3%<br>p = 0.8<br>slope = 0.01<br>F(1,20) = 0.06 | R<sup>2</sup> = 2%<br>p = 0.5<br>slope = 3.5<br>F(1,20) = 4.1 | R<sup>2</sup> = 13%<br>p = 0.09<br>slope = -0.7<br>F(1,20) = 3.1 |
|  | ALL | R<sup>2</sup> = 0.6%<br>p = 0.35<br>slope = -0.09<br>F(1,175) = 0.9 | R<sup>2</sup> = 3%<br>p = 0.04<br>slope = 0.03<br>F(1,175) = 4.2 | R<sup>2</sup> = 1%<br>p = 0.2<br>slope = 0.55<br>F(1,175) = 1.5 | R<sup>2</sup> = 0.2%<br>p = 0.55<br>slope = -0.08<br>F(1,175) = 0.3 |
|  | GC ad | R<sup>2</sup> = 4%<br>p = 0.24<br>slope = -0.4<br>F(1,33) = 1.4 | R<sup>2</sup> = 11%<br>p = 0.06<br>slope = -0.07<br>F(1,33) = 3.9 | R<sup>2</sup> = 17%<br>p = 0.015<br>slope = -4.3<br>F(1,33) = 6.6 | R<sup>2</sup> = 5%<br>p = 0.2<br>slope = -0.3<br>F(1,33) = 1.7 |
| Spiketrain Reliability (R<sub>w</sub>) | GC yo | R<sup>2</sup> = 2%<br>p = 0.2<br>slope = 0.001<br>F(1,100) = 1.8 | R<sup>2</sup> = 6%<br>p = 0.03<br>slope = 4e-3<br>F(1,100) = 4.7 | R<sup>2</sup> = 7%<br>p = 0.02<br>slope = 0.01<br>F(1,100) = 6.1 | R<sup>2</sup> = 3%<br>p = 0.17<br>slope = 0.002<br>F(1,100) = 1.9 |
|  | FS | R<sup>2</sup> = 48%<br>p = 0.0006, slope = 0.01<br>F(1,18) = 16.8 | R<sup>2</sup> = 70%<br>p < 0.0001<br>slope = -0.007<br>F(1,18) = 41.7 | R<sup>2</sup> = 6%<br>p = 0.29<br>slope = -0.12<br>F(1,18) = 0.7 | R<sup>2</sup> = 39%<br>p = 0.003<br>slope = -0.014<br>F(1,18) = 11.6 |
|  | HMC | R<sup>2</sup> = 29%<br>p = 0.19<br>slope = 0.005<br>F(1,16) = 6.5 | R<sup>2</sup> = 24%<br>p = 0.06<br>slope = -0.001<br>F(1,16) = 5.0 | R<sup>2</sup> = 9%<br>p = 0.2<br>slope = 0.006<br>F(1,16) = 0.3 | R<sup>2</sup> = 3%<br>p = 0.5<br>slope = -0.002<br>F(1,16) = 0.5 |
|  | CA3 +gzn | R<sup>2</sup> = 4%<br>p = 0.5<br>slope = 0.003<br>F(1,13) = 0.5 | R<sup>2</sup> = 9%<br>p = 0.29<br>slope = 6e-4<br>F(1,13) = 1.2 | R<sup>2</sup> = 4%<br>p = 0.45<br>slope = -0.01<br>F(1,13) = 0.45 | R<sup>2</sup> = 10%<br>p = 0.24<br>slope = -0.004<br>F(1,13) = 1.5 |
|  | GC +gzn | R<sup>2</sup> = 2%<br>p = 0.5<br>slope = -0.003<br>F(1,20) = 0.4 | R<sup>2</sup> = 22%<br>p = 0.03<br>slope = 8e-4<br>F(1,20) = 5.7 | R<sup>2</sup> = 17%<br>p = 0.05<br>slope = -0.02<br>F(1,20) = 3.9 | R<sup>2</sup> = 29%<br>p = 0.01<br>slope = 0.008<br>F(1,20) = 8.0 |
|  | ALL | R<sup>2</sup> = 0.4%<br>p = 0.5<br>slope = 7e-4<br>F(1,175) = 0.5 | R<sup>2</sup> = 7%<br>p = 0.0013<br>slope = -4e-4<br>F(1,175) = 10.7 | R<sup>2</sup> = 3%<br>p = 0.045<br>slope = -0.008<br>F(1,175) = 4.1 | R<sup>2</sup> = 1%<br>p = 0.24<br>slope = 0.001<br>F(1,175) = 1.4 |
|  | GC ad | R<sup>2</sup> = 13%<br>p = 0.03<br>slope = 0.006<br>F(1,33) = 4.9 | R<sup>2</sup> = 6%<br>p = 0.17<br>slope = 4e-4<br>F(1,33) = 2.0 | R<sup>2</sup> = 24%<br>p = 0.003<br>slope = 0.04<br>F(1,33) = 10.3 | R<sup>2</sup> = 8.5%<br>p = 0.09<br>slope = 0.004<br>F(1,33) = 3.05 |

**Table S2.** Spike-wise neural noise

| y-axis ↓ | x-axis → |  |  |  |
| --- | --- | --- | --- | --- |
|  |  | Delay | Jitter | Spike Probability (SP) |
| Normalized Decorrelation | GC yo | $R^2 = 20\%$<br>$p < 0.0001$<br>slope = 1.1<br>$F(1,100) = 24.5$ | $R^2 = 31\%$<br>$p < 0.0001$<br>slope = 1.05<br>$F(1,100) = 45.1$ | $R^2 = 41\%$<br>$p < 0.0001$<br>slope = -37.6<br>$F(1,100) = 69.8$ |
| | FS | $R^2 = 11\%$<br>$p = 0.15$<br>slope = 2.3<br>$F(1,18) = 2.2$ | $R^2 = 18\%$<br>$p = 0.06$<br>slope = 9.6<br>$F(1,18) = 3.9$ | $R^2 = 88\%$<br>$p < 0.0001$<br>slope = -49.1<br>$F(1,18) = 129.8$ |
| | HMC | $R^2 = 9\%$<br>$p = 0.22$<br>slope = 2.1<br>$F(1,16) = 1.6$ | $R^2 = 18\%$<br>$p = 0.08$<br>slope = 3.4<br>$F(1,16) = 3.5$ | $R^2 = 34\%$<br>$p = 0.011$<br>slope = -42.0<br>$F(1,16) = 8.1$ |
| | ALL | $R^2 = 35\%$<br>$p < 0.0001$<br>slope = 1.9<br>$F(1,175) = 74.3$ | $R^2 = 23\%$<br>$p < 0.0001$<br>slope = 1.4<br>$F(1,175) = 41.0$ | $R^2 = 57\%$<br>$p < 0.0001$<br>slope = -50.6<br>$F(1,175) = 183.8$ |
| | GC ad | $R^2 = 2\%$<br>$p = 0.38$<br>slope = 0.9<br>$F(1,33) = 0.8$ | $R^2 = 4\%$<br>$p = 0.23$<br>slope = -3.5<br>$F(1,33) = 1.4$ | $R^2 = 42\%$<br>$p < 0.0001$<br>slope = -45.7<br>$F(1,33) = 23.8$ |
| | Shuffle (GC yo) | $R^2 = 5\%$<br>$p = 0.02$<br>slope = 0.6<br>$F(1,100) = 5.8$ | $R^2 = 20\%$<br>$p < 0.0001$<br>slope = 0.8<br>$F(1,100) = 25.6$ | $R^2 = 61\%$<br>$p < 0.0001$<br>slope = -36.5<br>$F(1,100) = 160.0$ |
| Spiketrain Reliability ($R_w$) | GC yo | $R^2 = 23\%$<br>$p < 0.0001$<br>slope = -0.012<br>$F(1,100) = 29.9$ | $R^2 = 37\%$<br>$p < 0.0001$<br>slope = -0.012<br>$F(1,100) = 58.9$ | $R^2 = 48\%$<br>$p < 0.0001$<br>slope = 0.42<br>94.1 |
| | FS | $R^2 = 10\%$<br>$p = 0.16$<br>slope = -0.025<br>$F(1,18) = 2.1$ | $R^2 = 11\%$<br>$p = 0.15$<br>slope = -0.084<br>$F(1,18) = 2.2$ | $R^2 = 87\%$<br>$p < 0.0001$<br>slope = 0.55<br>$F(1,18) = 117.7$ |
| | HMC | $R^2 = 36\%$<br>$p = 0.009$<br>slope = -0.030<br>$F(1,16) = 8.8$ | $R^2 = 53\%$<br>$p = 0.0007$<br>slope = -0.043<br>$F(1,16) = 17.7$ | $R^2 = 31\%$<br>$p = 0.016$<br>slope = 0.30<br>$F(1,16) = 7.3$ |
| | ALL | $R^2 = 40\%$<br>$p < 0.0001$<br>slope = -0.020<br>$F(1,175) = 93.4$ | $R^2 = 28\%$<br>$p < 0.0001$<br>slope = -0.016<br>$F(1,175) = 53.4$ | $R^2 = 61\%$<br>$p < 0.0001$<br>slope = 0.5<br>$F(1,175) = 219.1$ |
| | GC ad | $R^2 = 7\%$<br>$p = 0.14$<br>slope = -0.01<br>$F(1,33) = 2.3$ | $R^2 = 0.03\%$<br>$p = 0.9$<br>slope = 0.003<br>$F(1,33) = 0.01$ | $R^2 = 16\%$<br>$p = 0.017$<br>slope = 0.2<br>$F(1,33) = 6.3$ |
| | Shuffle (GC yo) | $R^2 = 5\%$<br>$p = 0.03$<br>slope = -0.006<br>$F(1,100) = 5.0$ | $R^2 = 21\%$<br>$p < 0.0001$<br>slope = -0.008<br>$F(1,100) = 26.2$ | $R^2 = 62\%$<br>$p < 0.0001$<br>slope = 0.4<br>$F(1,100) = 162$ |

**Table S3.** Spiketrain-wise properties

| y-axis ↓<br>x-axis → | | Overall Firing Rate | Spiketrain Reliability ($R_w$) |
| --- | --- | --- | --- |
| Normalized Decorrelation | GC<br>yo | $R^2 = 15\%$<br>$p < 0.0001$<br>slope = - 1.3<br>$F(1,100) = 18.4$ | $R^2 = 81\%$<br>$p < 0.0001$<br>slope = -87<br>$F(1,100) = 430.1$ |
| | FS | $R^2 = 65\%$<br>$p < 0.0001$<br>slope = - 0.5<br>$F(1,18) = 33.5$ | $R^2 = 90\%$<br>$p < 0.0001$<br>slope = -85<br>$F(1,18) = 169.0$ |
| | HMC | $R^2 = 35\%$<br>$p = 0.0010$<br>slope = - 2.8<br>$F(1,16) = 8.6$ | $R^2 = 61\%$<br>$p = 0.0001$<br>slope = -105<br>$F(1,16) = 25.5$ |
| | CA3<br>+gzn | $R^2 = 69\%$<br>$p = 0.0001$<br>slope = - 1.7<br>$F(1,13) = 28.7$ | $R^2 = 55\%$<br>$p = 0.0016$<br>slope = -76<br>$F(1,13) = 15.8$ |
| | GC<br>+gzn | $R^2 = 0.6\%$<br>$p = 0.73$<br>slope = - 0.2<br>$F(1,20) = 0.1$ | $R^2 = 28\%$<br>$p = 0.0111$<br>slope = -75<br>$F(1,20) = 7.8$ |
| | ALL | $R^2 = 37\%$<br>$p < 0.0001$<br>slope = - 0.8<br>$F(1,175) = 101.5$ | $R^2 = 79\%$<br>$p < 0.0001$<br>slope = -91<br>$F(1,175) = 673.1$ |
| | GC<br>ad | $R^2 = 30\%$<br>$p = 0.0006$<br>slope = - 4.1<br>$F(1,33) = 14.5$ | $R^2 = 52\%$<br>$p < 0.0001$<br>slope = -90<br>$F(1,33) = 36.2$ |

**Table S4.**
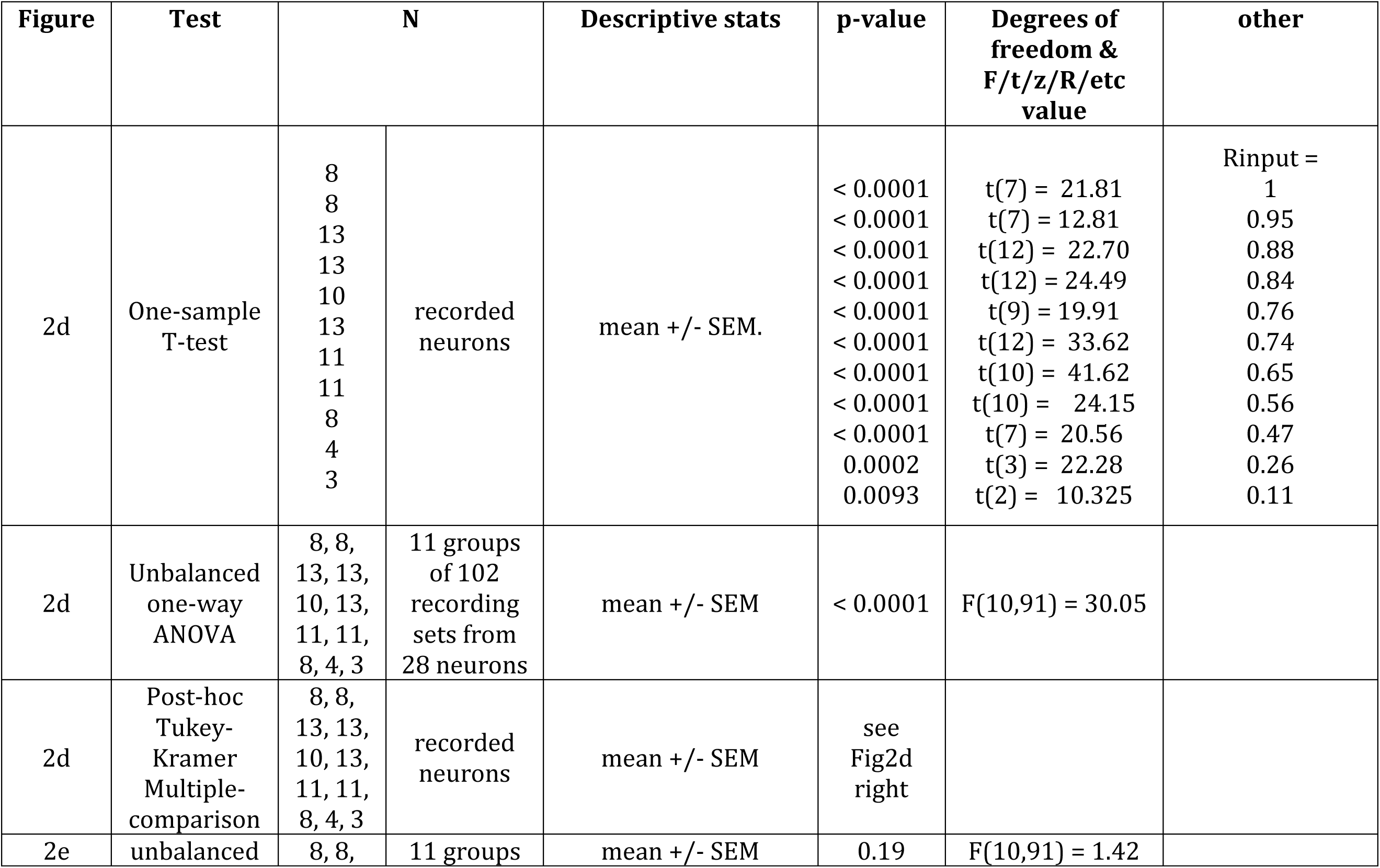
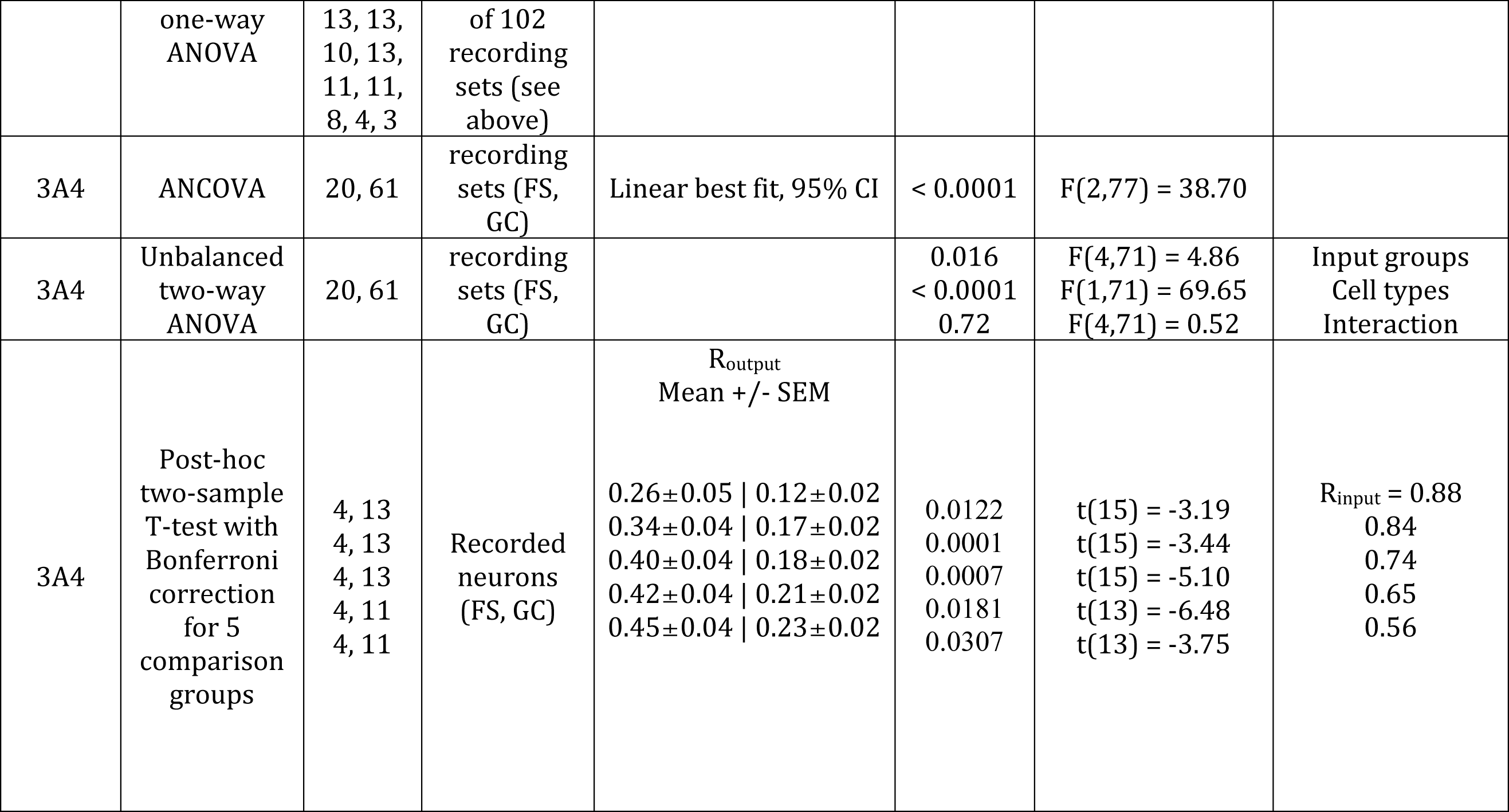
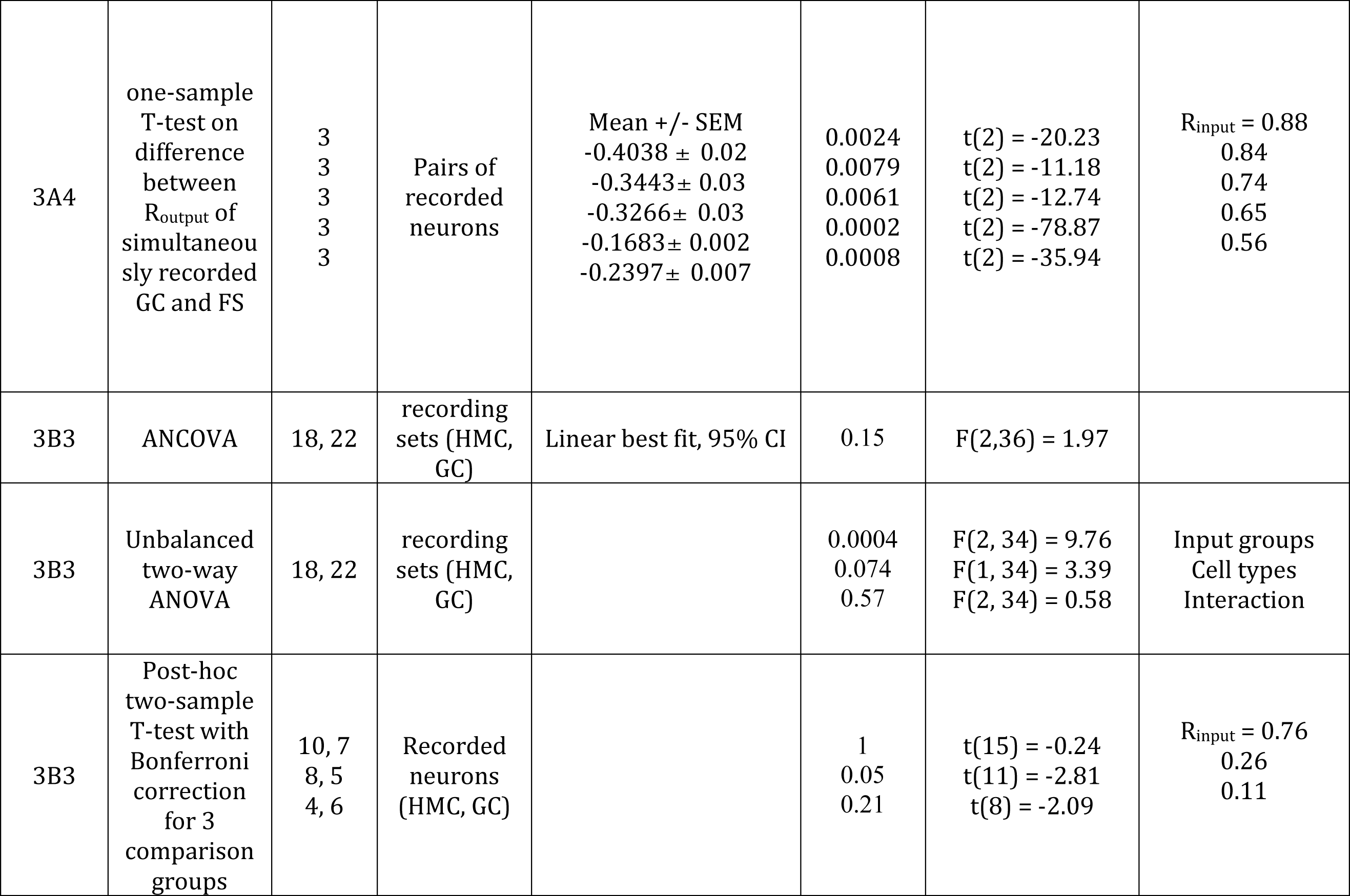
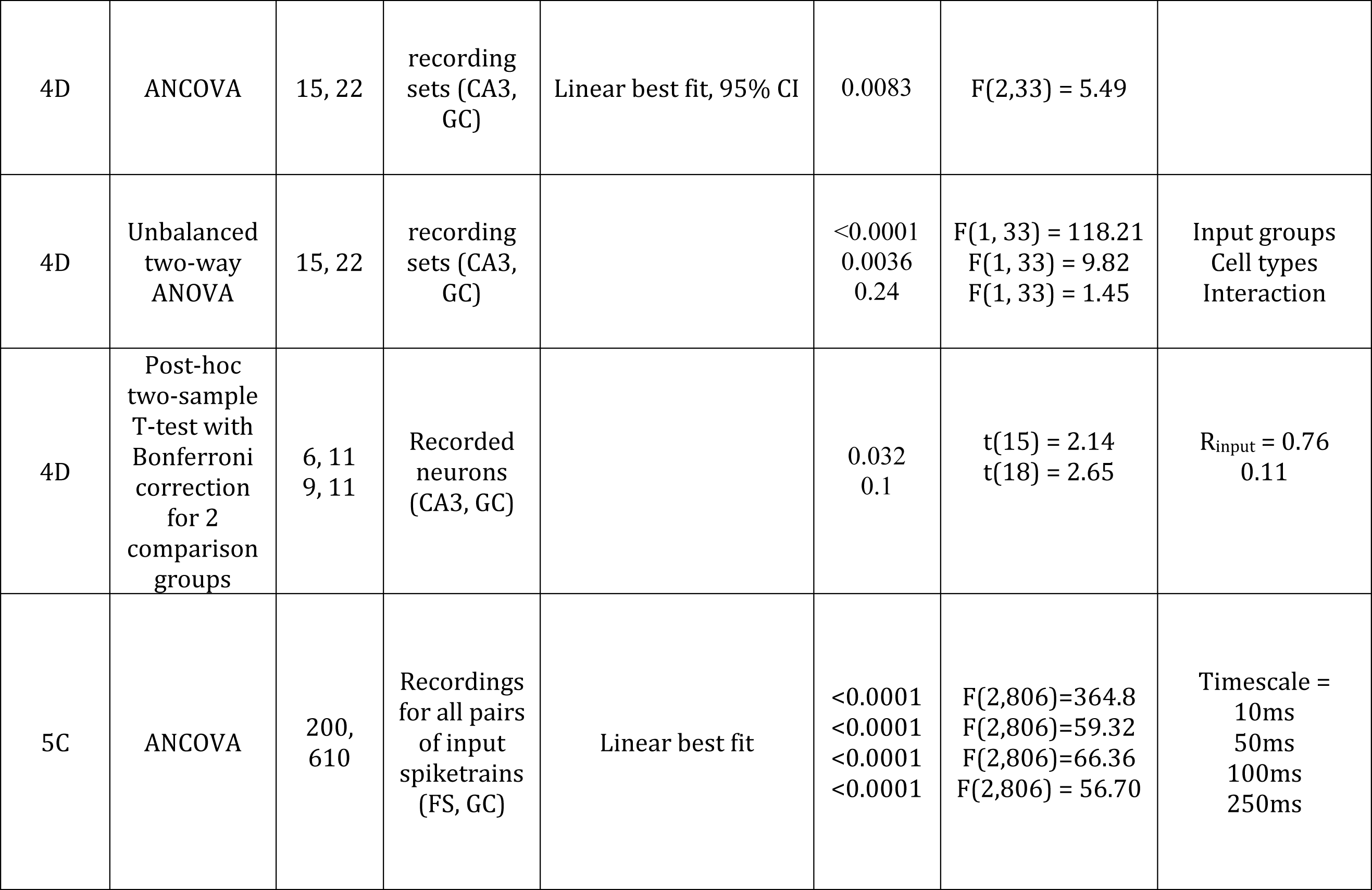
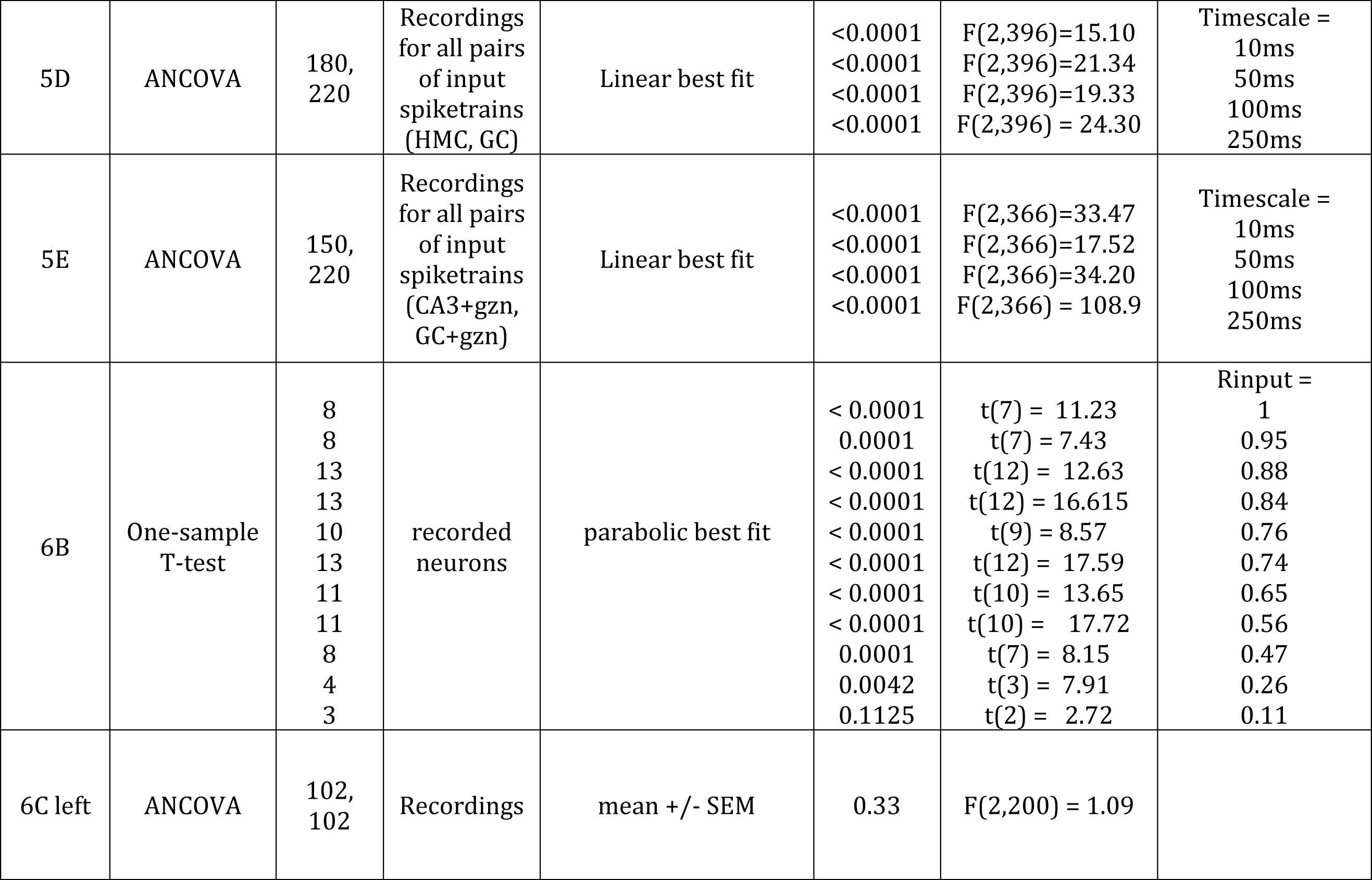
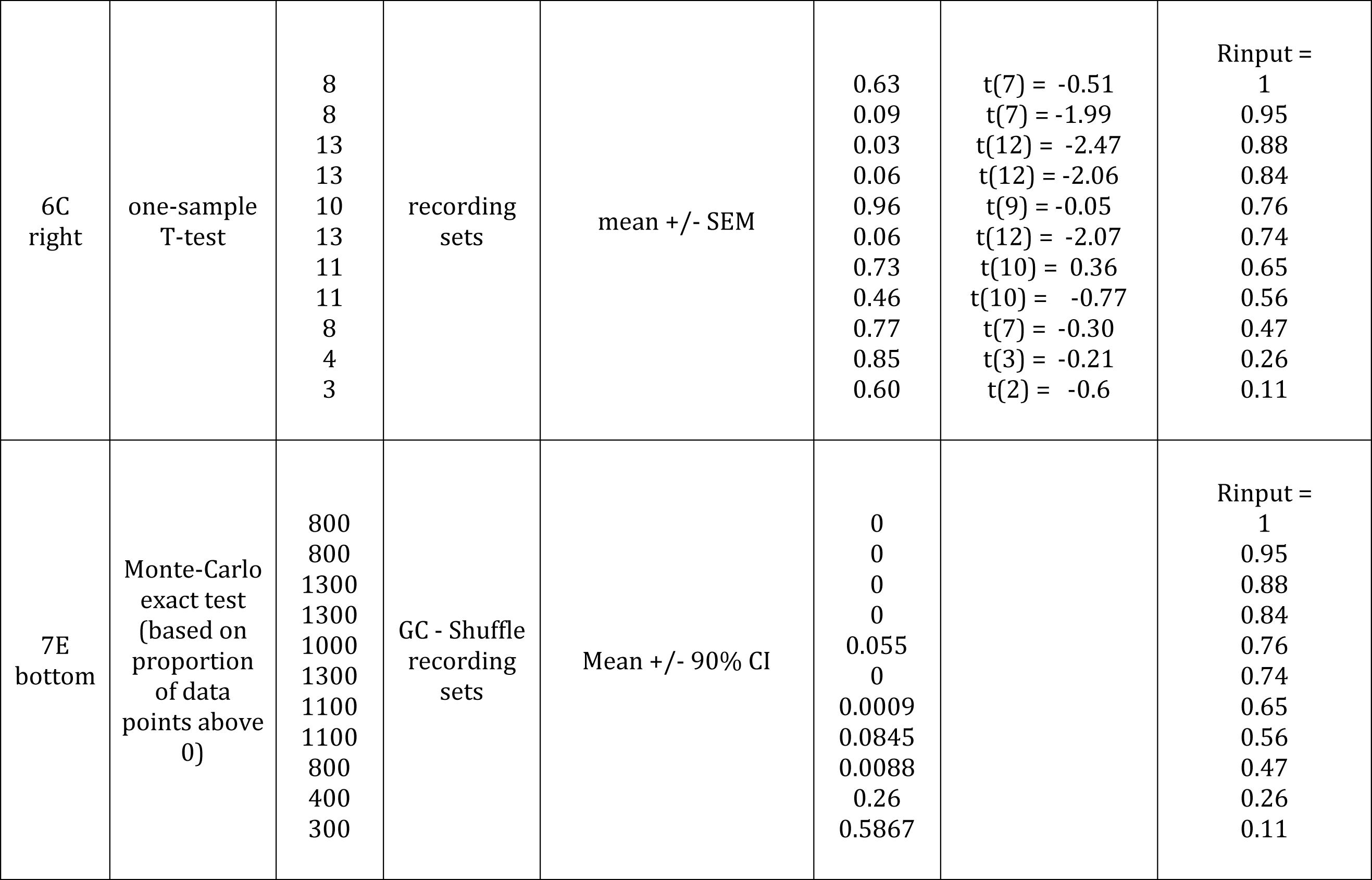
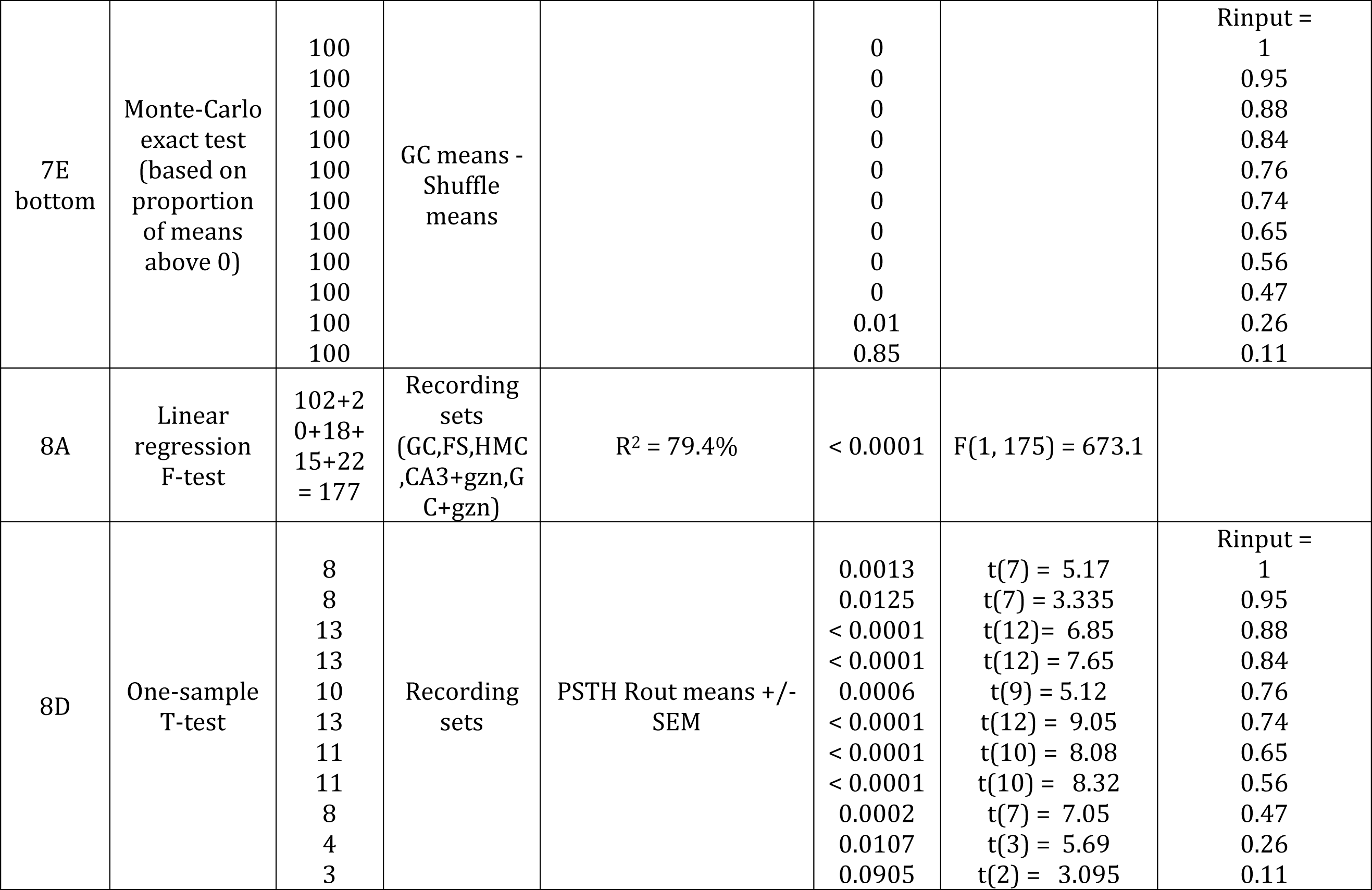
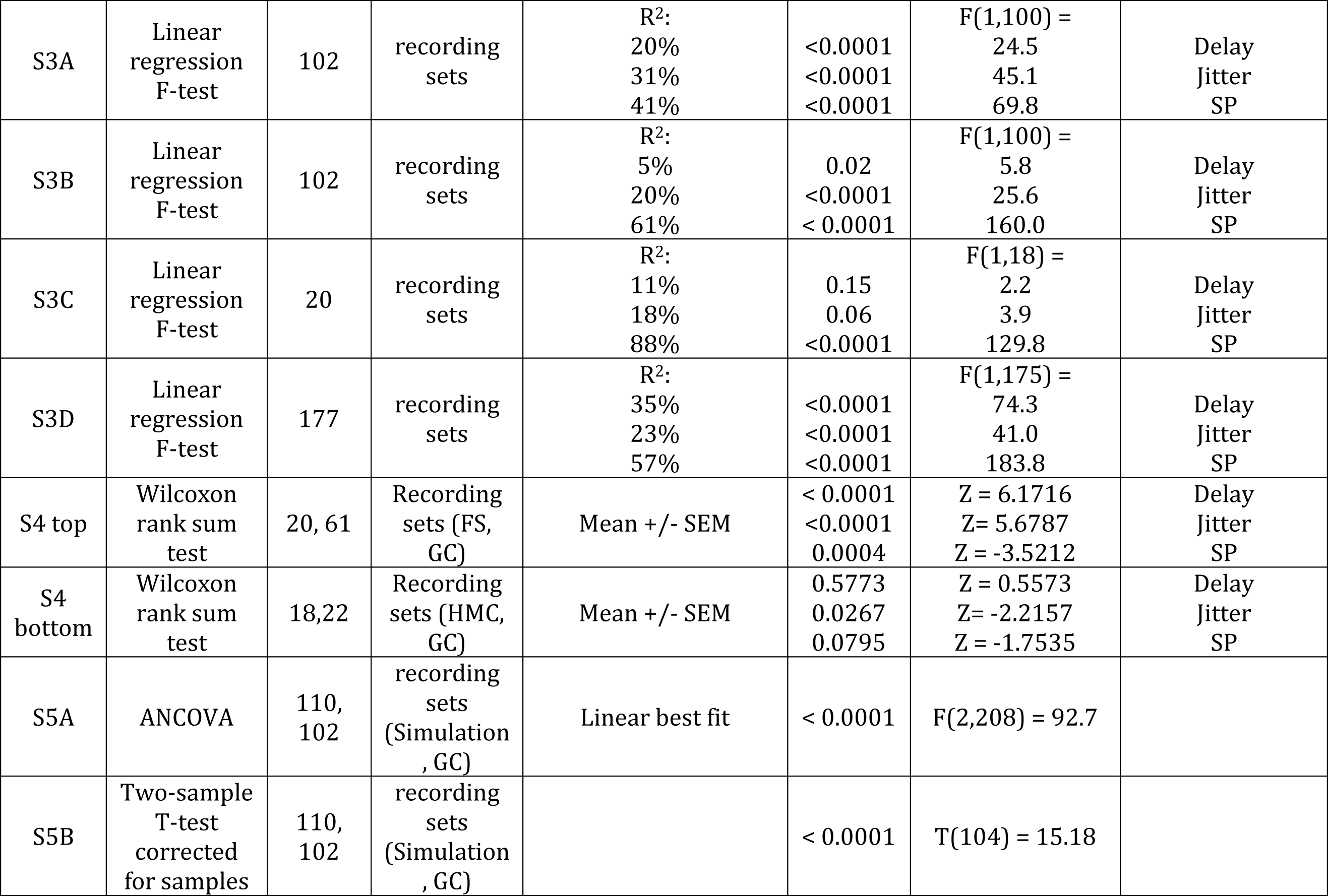
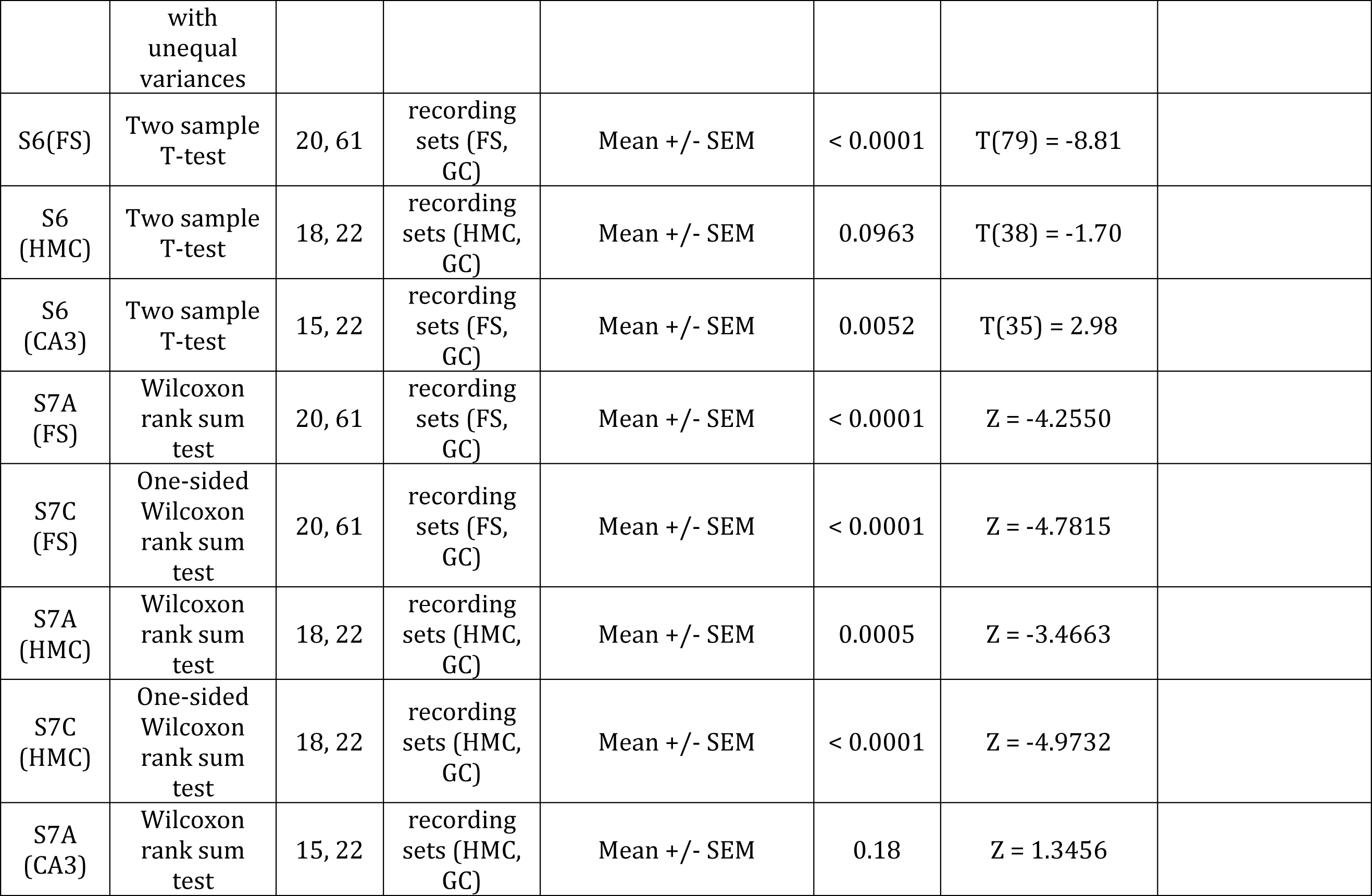
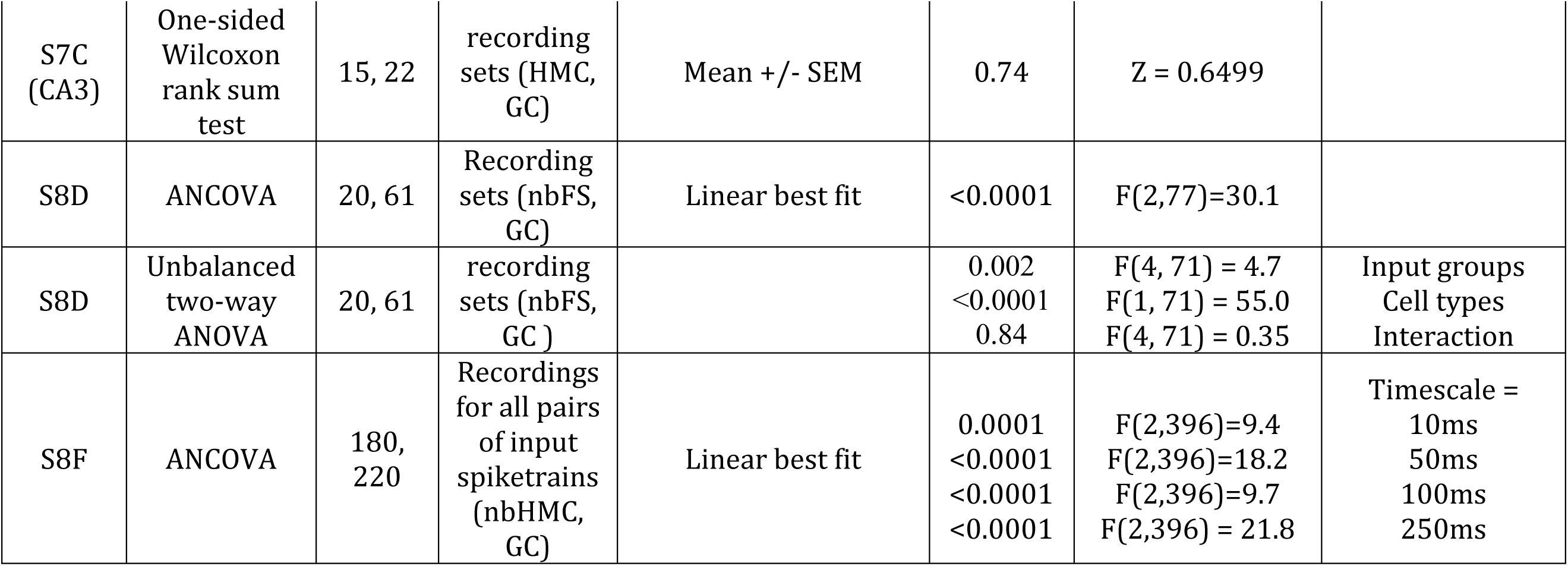
Statistics

## Acknowledgements

We would like to thank Jesse Pfammatter and Linda Overstreet-Wadiche for many helpful discussions and feedback on early versions of the manuscript.

